# Optimizing Niosome Formulations for Enhanced Cellular Applications: A Comparative Case Study with L-α-lecithin Liposomes

**DOI:** 10.1101/2023.11.14.567080

**Authors:** Nilufer Cakir, Naile Ozturk, Asli Kara, Ali Zarrabi, Nur Mustafaoglu

## Abstract

This study delves into the optimization of niosome production for biological applications, focusing on their emerging role as amphiphilic nanoparticles derived from nonionic surfactants, poised at the forefront of biomedical research. We aimed to formulate and characterize a diverse array of niosomal nanoparticles, with particular emphasis on process-related parameters and physicochemical characteristics. Critical thresholds for size, polydispersity, and zeta potential were established to identify parameters crucial for optimal niosomal formulations through a comprehensive investigation of concentrations, sonication times, ingredient ratios, and surfactant types. Leveraging MODDE® software, we generated ten optimized formulations from preliminary parameter screening. The proposed experimental model design by the software exhibited acceptable similarity to the obtained experimental results (F-score:0.83). The criteria for selection of the predicted experimental model formed based on targeted physicochemical considerations. To enhance half-life and penetration, especially in higher electrostatic regions like the Central Nervous System (CNS), we proposed a neutralized surface charge (−10 to 10 mV) while maintaining size within 100-200 nm and polydispersity below 0.5.

Extended stability screening revealed periodic and extended Gaussian distributions for size and zeta potential to minimize flocculation and coagulation caused by neutralized surface charge. Notably, the cellular response performance of optimized niosomes was assessed via cellular binding, uptake, and viability in comparison to liposomes. Glioblastoma cell line (U-87) and granulocyte colony-stimulating factor (G-CSF) containing lymphoblastic leukemia cell line (NFS-60) were chosen to represent tumors developed in the CNS region and white blood cells, respectively, enabling a comprehensive comparative analysis with liposomes.

The meticulous comparison between niosomes and liposomes revealed comparable cellular viability profiles on both U-87 and NFS-60 cell lines, highlighting their similarities in cellular interactions. Moreover, selected niosomal formulations demonstrated exceptional cellular uptake, either equaling or surpassing observed liposomal uptake. One of the most promising niosomes was selected and optimized to evaluate drug encapsulation performance of niosomes for further drug delivery adaptations by one of chemotherapy drugs, Paclitaxel (PTX). Cytotoxicity study was established with the most efficiently encapsulated niosome condition with human-derived fibroblasts (HDFs) and U-87 as the representation of healthy and cancerous cell lines. Results demonstrated 1:100 diluted PTX-loaded niosome in the certain concentration demonstrated favourable toxicity in U-87 than original PTX at the same concentration while not disturbing healthy HDFs. These findings underscore the potential of niosomes for reliable drug delivery, challenging the dominance of liposomal vehicles and presenting economically viable nanocarriers with significant implications for advancing biomedical research.

## 1. Introduction

Niosomes are one of the promising pseudo-lipid nanostructures for the delivery of natural agents and drugs, which have gained acceptance as structural analogs of liposomes in the development of drug delivery strategies. Niosomes have become particularly popular for drug delivery in recent years due to their amphiphilic, biocompatible, biodegradable, and nontoxic properties. Unlike liposomes, niosomes have a vesicle construct that contains nonionic surfactants and forms a bilayer that provides better bioavailability by increasing residence time and decreasing renal clearance.^(1)–(4)^ Niosomes are useful not only for drug delivery but also for targeted drug transport mechanisms such as antigens and small molecules.^(5)^ Due to the use of nonionic surfactants, they are a cost-effective alternative; their longer chemical stability also makes them a promising candidate for drug delivery applications.^(1),(4),(6),(7)^ They can encapsulate the drug into vesicles that improve drug bioavailability, therapeutic efficiency, and penetration through the tissue, release the drug in a sustained and controlled manner, and can be targeted to the desired site by adjusting the composition, which helps to reduce side effects.^(8)^

Advances of niosomes are not sufficient to develop cost-effective alternatives to liposomes. The cellular application gap between niosome and liposome are needed to be filled and well-understood in the literature. Several studies have demonstrated the advantages of lipid nanoparticles as smart drug carriers through active or passive targeting.^(9)–(11)^ In parallel, there are niosomal nanoparticles designed more cost-effectively and demonstrated improved encapsulation yields, resulting stable, and reproducible nanocarrier formulations.^(12)^ However, simultaneous comparison with liposomes and exact cellular application performance were not reported to demonstrated the cost-effective alternative of liposomes. Additionally, the influence of surfactants on the physical properties of niosomal nanoparticles has been explored as enhancer toward the stability compared to lipid nanoparticles. Nevertheless, most studies have focused on drug-loaded lipid or pseudolipid nanoparticles, particularly *in vitro* testing and drug-release profiling, while little attention has been paid to the properties and characteristics of niosomal nanoparticle formulations and cellular performance differences/similarities with liposomes that can be effectively used in drug delivery purposes.

The critical process attributes for reproducing niosomal nanoparticles and achieving appropriate cellular interactions with them are main considerations, specially from the drug delivery framework.^(2),(3),(13)–(16)^ Thus, while this study may lead to effective drug delivery profiles, the potential alternatives to liposomes may literally be recorded based on their simultaneous performance. Therefore, the main objective of this study is to optimize the physicochemical and biological properties of drug-free niosomal nanoparticles, which have not been extensively studied in the literature with preliminary set criteria by the concern of crossing blood-brain barrier (BBB). Niosomes have been investigated for various drug delivery mechanisms, such as targeted and controlled drug delivery across the blood-brain barrier (temozolomide)^(13)^, the eyes (tacrolimus, naltrexone HCl)^(17)^ transdermally (gallidermin, clopramine)^(18)^, pulmonarily (glucocorticoid)^(19)^ and orally (cefdinir, lornoxicam)^(2)^ Synchronous delivery of anticancer drugs via niosomes has been used for doxorubicin and curcumin.^(20),(21)^

The design of excipients for drug development requires compliance with several commitments, including optimization of the nanostructure formulation with respect to factors such as penetration, minimum effective concentration, minimum toxic concentration, bioavailability at the site of action, frequency and route of administration, and physicochemical state.^(22)^ Because of obligations, the carrier formulation and its optimization are as important as the drug itself.^(22)^ To achieve the optimal drug carrier formulation, it is critical to characterize the parameters related to nanoparticle geometry^(23)^, physicochemical properties^(6),(7),(24),(25)^, cellular toxicity^(24),(26)^, cellular uptake^(27),(28)^, and species-specific abundances^(29),(30)^. Few studies have used formulation-dependent optimization strategies, yet, niosomal formulations can be quite critical in biological applications, as demonstrated by the example of surfactant composition and surface charge of niosomes, which have a tremendous impact on the oral adsorption of repaglinide.^(31)^ On the other hand, the influence of surfactant type and production parameters such as concentration, sonication time, temperature, and surfactant/cholesterol ratio on drug delivery has not been fully elucidated, simultaneously.^(3),(32),(33)^ There is ample evidence in the literature that niosomes are an alternative to liposomes in terms of efficacy, safety, and stability, but the lack of reliable and reproducible niosomal formulations remains still as a limitation for transiting of these literature findings into the clinic.^(34)–(36)^ Therefore, it is necessary to conduct further characterization studies of niosomal formulations to optimize the process parameters for their reliable and reproducible production.

Here we emphasize the importance of preliminary analytical characterization of drug carriers, niosomes, and propose several niosome formulation alternatives with critical process parameter evaluation. Overall, preliminary physicochemical characterizations, cellular toxicity, cellular uptake, drug encapsulation performance and formulation-specific frequencies to address carrier-mediated critical parameters and simultaneous comparison with liposome nanostructures from the same analytical perspective provide evidence to the literature on the compatibility of niosomes, with the goal of more cost-effective production approach than liposomes for drug delivery, not only for research-based studies but also for pharmaceutical industry. *Figure 1* shows the general outline of the study and clarifies the perspective of the study.

**Figure 1.**
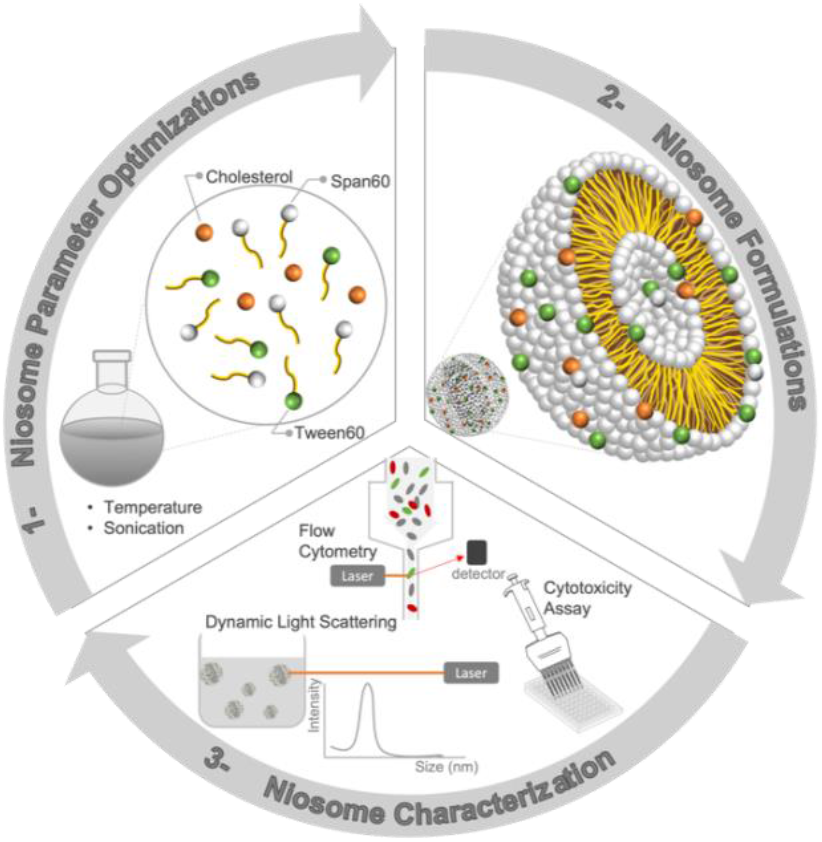
Overall study perspective for determining quality attributes and critical process parameters of niosomes used in drug delivery system.

## 2. Materials and Methods

### Materials

Span 60 (Sigma Aldrich, Germany), Tween 80 (Sigma Aldrich, Germany), fluorescein isothiocyanate (FITC) (Sigma Aldrich, Germany), L-α-lecithin, and Cholesterol (Chol) (Sigma Aldrich, Germany) were purchased from Sigma-Aldrich Chemical Co. Chloroform was purchased from Merck (Germany). 3-(4,5-Dimethylthiazol-2-yl)-2,5-diphenyltetrazolium bromide (MTT) was purchased from Merck (Germany). The U-87 cell line was a kind gift from Esendagli Group at Hacettepe University (Turkey). The NFS-60 cell line was a gift from ILKOGEN Biotech. All organic solvents were analytical grade and deionized distilled water was obtained from Merck Millipore with a fixed 18.2-ohm conductivity.

### Preparation of niosomes ^(37)^

Various niosome formulations (*Table 1*) containing different types of surfactants and molar ratios were prepared using the thin-layer hydration technique. ^(34),(38),(39)^ The appropriate amount of cholesterol and surfactants were dissolved in a mixture of ethanol and chloroform (0.1:1 v/v) in a round bottom flask with a final volume of 50 mL. The formulations are classified as multiple surfactant type formulations (MSTF) and single surfactant type formulations (SSTF). 3 different molar ratios for MSTF and 2 types of surfactants for SSTF in a molar ratio of 1:1 with cholesterol were established as preliminary niosome characterizations. A total of five different formulations were formed and designated as Formulation A, Formulation B, Formulation C, Formulation D and Formulation E. Four different final solution concentrations were set as final solution concentration and composition (%) in mass was considered for ingredients by considering the final solution concentrations. Two different sonication times, 40 and 90 minutes, were set and applied for all five types of formulations. All preliminary experimental conditions were represented in *Table 1*. Each formulation was prepared in triplicate, at four different concentrations, and with two different sonication times; therefore, 24 samples were prepared for each formulation, resulting in a total of 120 samples. The organic phase was removed at 60°C under vacuum using a rotary evaporator (Heidolph, Germany). The vacuum time was set at 1 hour for all formulations, and the dried thin film was hydrated with 10 mL of phosphate buffer saline (PBS) (Sigma-Aldrich Co., Germany) at 60°C by gently shaking the round bottom flask until all the dried film was dissolved. Each prepared formulation was treated in a sonication bath (ISOLAB, Turkiye) for 45 and 90 minutes, respectively. In this study, sonication time is identified as a critical factor in controlling nanoparticle size, with an optimal range minimizing particle size. Insufficient or excessive sonication can lead to suboptimal outcomes, such as increased particle size or potential nanoparticle degradation. Therefore, sonication times above and below the reference value of 60 minutes were evaluated to determine the optimal duration for achieving the desired nanoparticle size range in nanometers.

**Table 1.**
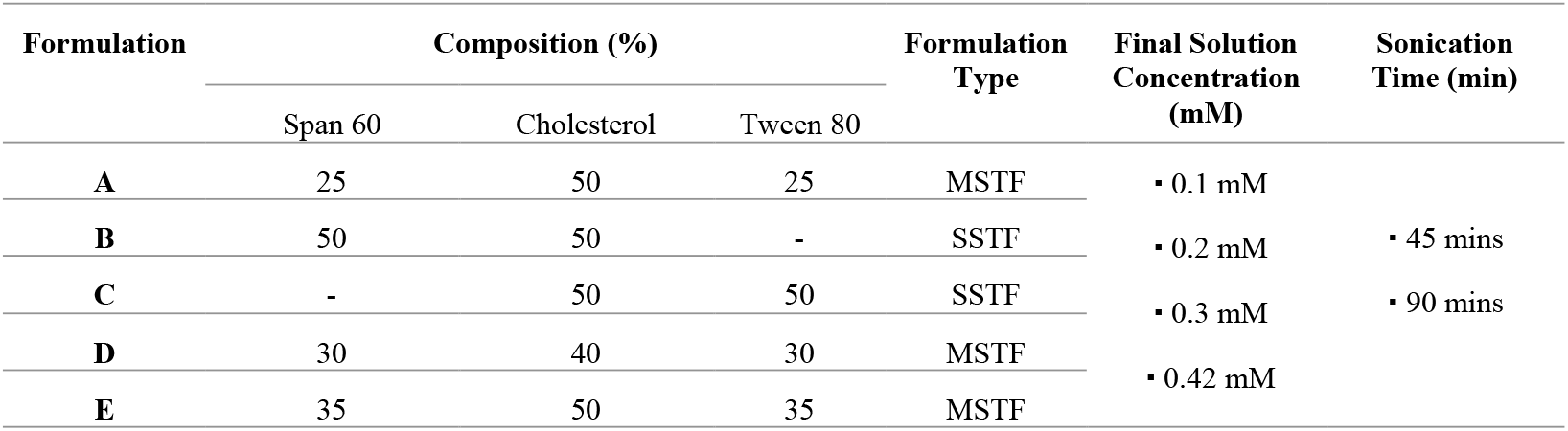
Preliminary screened formulation A, B, C, D, and E ingredient and condition list.

### Preparation of niosomal nanoparticles ^(33,37)^

The experimental process began with careful preparation, combining surfactants and cholesterol within round-bottom bottles, dissolved in chloroform to achieve a consistent liquid. Utilizing a rotary evaporator under controlled conditions at 60°C and adjusted vacuum settings resulted in the formation of a thin layer at the bottle’s base. Subsequent hydration using Phosphate Buffered Saline (PBS) (Sigma-Aldrich, Germany) followed by ultrasonic treatment generated milky-white pseudo-lipid nanoparticles.

Fine-tuning parameters of niosome production were originated via thin-film hydration method followed by sonication for achieving desired size and homogeneity. Sonicated mixtures at the determined conditions were filtered via several membrane filtering and transferred to amber glass vials for long-term preservation at 2-8 °C. Pore size of membrane filters was 200 μm and 100 μm respectively, may require additional filtering depends on final mixture concentration. The synthesis aimed for nanoparticles ranging from 100-200 nm with homogeneity between 0.2 and 0.5, employing a ‘thin-film layer’ technique^(33)^, explicated in *Figure 2*.

**Figure 2.**
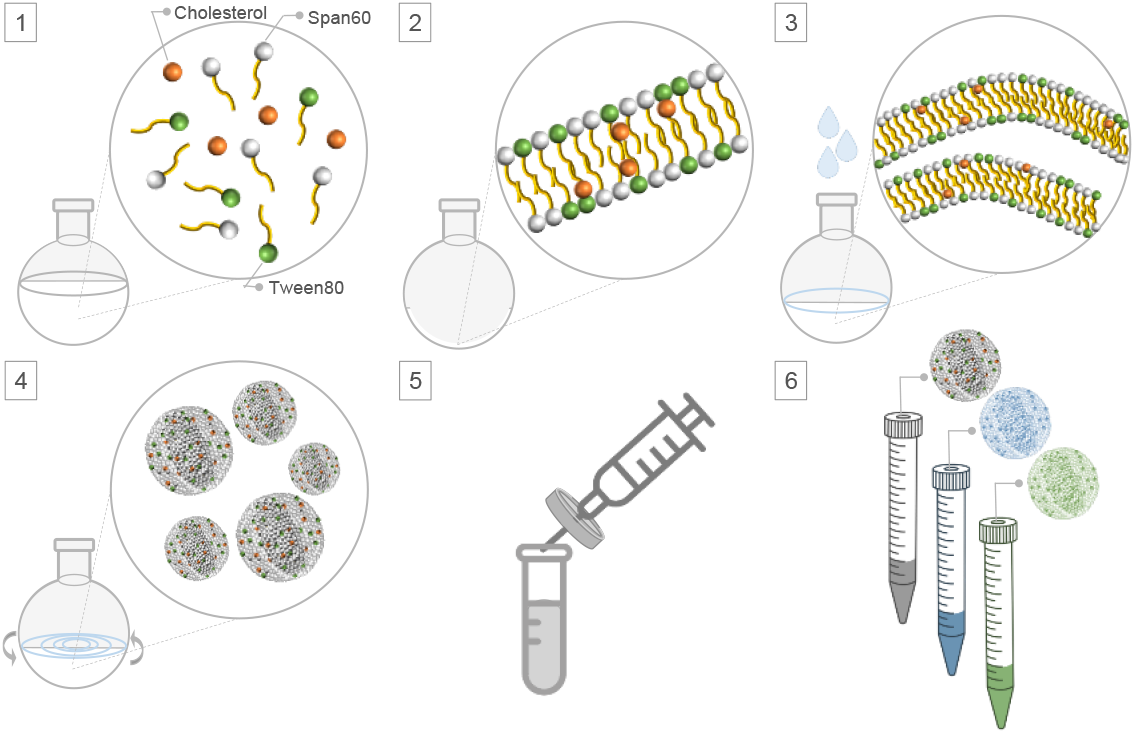
Thin-film layer method for preparation of niosomes: 1) Surfactant and cholesterol mixture, 2) Evaporation of organic solvent, 3) Hydration 4) Hydration with sonication, 5) Sterile filtration under the fume hood for further characterizations. 6) The collection of final niosomal nanoparticles.

The methodology involved precise initial composition preparation, the use of a rotary evaporator for controlled dissolution, hydration with PBS, and sonication for nanoparticle synthesis. Subsequent optimization, filtration, and comprehensive analysis through Dynamic Light Scattering (DLS) (Malvern, United Kingdom) facilitated a comprehensive assessment of homogeneity, size distribution, and surface charge.

### Preparation of liposomal nanoparticles:^(33)^

Liposomes were prepared by thin film hydration method.^(38)^ L-α-lecithin (3.3 mg/mL) and cholesterol (0.8 mg/mL) were dissolved in chloroform. The lipid mixture in chloroform solution was placed in a round bottom flask and the chloroform was evaporated using rotary evaporator (150 rpm, 55°C, 332 mbar) to form a lipid film on the sides of the round bottom flask. The resulting dry lipid film was hydrated with 3 mL of phosphate buffered saline (pH 7.4). For extrusion, the resulting lipid vesicles were passed through polycarbonate membranes of 0.6 μm, 0.4 μm and 0.2 μm pore sizes, respectively which is different than niosomal uniformity. Liposomes were then placed in a dialysis bag (3500 Da MWCO, regenerated cellulose membrane, Sigma Aldrich, Germany) and dialyzed against phosphate buffered saline (pH 7.4) for 16 hours. Size, polydispersity, and zeta potential measurement data for liposomes were given in the *Supplementary Table 6*.

### DLS Measurements

Physicochemical characterization was obtained by Dynamic-Light Scattering (DLS) (Malvern PanAnalytical, United Kingdom) with fixed and optimized experimental procedure. Briefly, each formulation was prepared as triplicates and 13 iterations were arranged for each run. 1:10 diluted PBS at pH:7.4 was assigned as blank before each triplicate to zeroize the system. Viscosity of niosomes was set as 1.48 ± 0.01 (mPa.s).^(39)^ To measure the size and homogeneity of formulations, polystyrene cuvettes (Malvern PanAnalytical, United Kingdom) were used. To measure zeta potential of each formulation, capillary zeta cell cuvettes (Malvern PanAnalytical, United Kingdom) were used.

### Fluorescence labelling of niosomes

Niosomes were labelled with fluorescein isothiocyanate (FITC) (Sigma-Aldrich, Germany) dissolved in ethanol at the fixed concentration and incubated overnight after rotary evaporation following the previously published thin-film hydration protocol.^(40),(41)^ Each niosome formulation is tagged by 300 nM FITC stock solution with two different mixing ratios (1:50 and 1:100 v/v). FITC entrapped inside of niosomes and excessive FITC was removed by continuous sterile filtering. EE (%) of FITC was calculated based on the former FITC-load absorbance and released FITC absorbance after centrifugal removal by using the equation demonstrated as *Eq*.*1*. The FITC concentration was calculated based on FITC-based standard curve demonstrated in *Supplementary Figure 6*. Each labelled formulation was physiochemically characterized after the labeling procedure by DLS with a dilution factor of 1:10 in PBS to determine whether its behavior changed in terms of size, charge, and polydispersity. The tagging of niosome-FITC was only generated for cellular uptake purposes. Calculated EE% for tagged niosomal formulations demonstrated in *Supplementary Figure 11*.

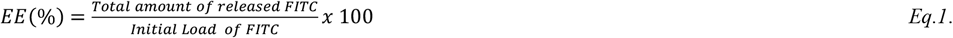

### Fluorescence labelling of liposomes

The liposomes were labeled with FITC using freeze-thaw cycles. ^(19),(42),(43)^ The method of freeze-thaw was accomplished after formation of liposomes. Thus, freeze-thaw cycles with the same amount of FITC were utilized to liposome followed by using extruder with 200 μm polycarbonate filtering. This involved subjecting the labeled liposomes to six subsequent freezing and thawing cycles, each cycle occurring every 4 hours. These cycles may induce stress on the liposomal membrane due to ice crystal formation during freezing.^(42)^ Therefore, FITC-tagged liposomes were checked via DLS after extruding and before treating with the cells to verify the preset required physicochemical specifications. Any excess FITC post freeze-thaw cycles were eliminated through 0.2 μm membrane filtration. Subsequent tracking of the size and homogeneity of the tagged liposomes allowed for simultaneous comparison with tagged niosomal nanoparticles, revealing expected levels of size, PDI, and zeta potential. Additional details can be found in *Supplementary Table 6*.

### Drug loading of niosomes

Paclitaxel (ATAXIL ®, DEVA Pharmaceuticals) was encapsulated by niosomes to determine their potential drug loading capacity (DLC%) and drug loading efficiency (DLE%) for future studies. Paclitaxel (PTX) loaded niosomes were then exposed to healthy (human-derived fibroblasts (HDFs) and cancerous cell lines (U-87) to investigate their potential cytotoxicity profiles. By this purpose, one of the most promising niosomes, Opt-10, was selected for exposure. Three different loading concentrations were prepared to load the niosome and the highest EE% corresponded concentration was selected for the toxicity studies. Required calculations were represented in *Eq*.*2, Eq*.*3*, and *Eq*.*4*. PTX standard curve was established with six different concentrations to determine EE% indicated in *Eq*.*1*. and DLE% demonstrated in *Eq*.*4*. All nanodrop measurements were performed at 230 nm which was established wavelength to determine PTX. ^(55)^ In all drug-loaded niosomes were analyzed by DLS before toxicity assay and results demonstrated in *Supplementary Figure 12*.*a* and *12*.*b*.

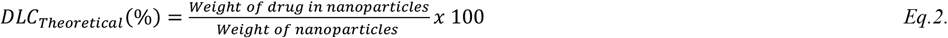

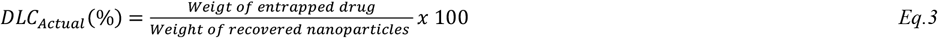

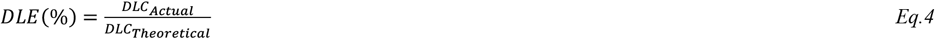

### FTIR analysis

FTIR analysis was performed to verify the presence of FITC in the niosomal nanoparticles and to ensure the labeling procedure. The samples were dried by freeze-drying in glass containers. The control groups consisted of FITC only and nanoparticles only to compare whether the labeling was successful. The dried samples were analyzed by FTIR (ThermoFisher, USA) and analyzed using OMNIC^®^ software. The statistical evaluations and the raw version of the data analyzed with the software are given in the *Supplementary Figure 8*.

### MODDE analysis

*P*reliminary formulation screening for investigation of final concentration, sonication time, surfactant type, and surfactant composition was evaluated by *MODDE*^*®*^ Software. Design model was set based on target size range as 100 – 200 nm. Modeling was determined as multifactorial analysis and significance was set as alpha: 0.5. Contour mapping mode and screening module were selected to obtain optimized formulations and 10 different optimization formulations were obtained. *Table 2* indicates *MODDE*^*®*^ Software output evaluated in logarithmic threshold.

**Table 2.**
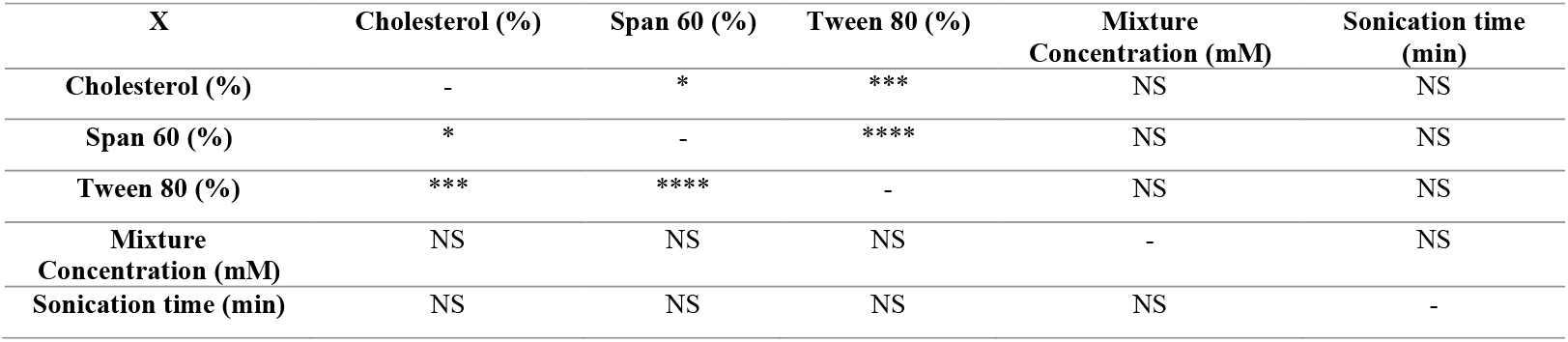
Two-factorial effect matrix of determined niosome production process parameters. (NS: p-value > 0.05; *: p-value ≤ 0.05; **: p-value ≤ 0.01; ***: p-value ≤ 0.001, ****: p-value ≤ 0.0001.)

| X | Cholesterol (%) | Span 60 (%) | Tween 80 (%) | Mixture Concentration (mM) | Sonication time (min) |
| --- | --- | --- | --- | --- | --- |
| Cholesterol (%) | - | * | *** | NS | NS |
| Span 60 (%) | * | - | **** | NS | NS |
| Tween 80 (%) | *** | **** | - | NS | NS |
| Mixture Concentration (mM) | NS | NS | NS | - | NS |
| Sonication time (min) | NS | NS | NS | NS | - |

### Cell maintenance

Glioblastoma cell line U-87 and lymphoblast cell line NFS-60 were grown as adherent and suspension cells in Dulbecco’s Modified Eagle Medium (DMEM) (Pan Biotech, Germany) containing 10% FBS + 1% Penicillin/Streptavidin and RPMI 1640 (Pan Biotech, Germany) + 10% M-CSF (Gibco), respectively. Cells were cultured in T75 culture flasks (ISOLAB Co, Germany.) at the same period (3-day culture interval) with a division ratio of 1:5.

### Cell viability assay

All cell viability assays were performed in 96-well plates (ISOLAB Co.) by seeding 5000 cells/well in 100 μL to each well with 2 hours incubation at 37°C. Except PTX loaded niosomes and their toxicity profile, all toxicity studies with non-load niosomes were performed with 9 serial dilutions with 1:10 dilution from 1000 pM to 10^−5^ pM. All dilutions’ first stock was prepared in starvation media with 1:2 volume ratio. Niosomal nanoparticles were incubated with the cells 24, 48, or 72 hours. Then, 3-(4,5-Dimethylthiazol-2-yl)-2,5-Diphenyltetrazolium Bromide (MTT) was dissolved in PBS at pH: 7.4 with a final concentration of 0.5 mg/mL, applied 50 μL on each well of the plated, incubated 3 hours. Reading of the wells was performed by removing media and adding 200 μL solubilization solution containing 10% SDS in DMF at pH:4.6. The results of cell viability assays were obtained using a microplate reader, detected at the wavelength of 570 nm. (BioRad, USA).

In PTX loaded niosomes, the highest EE% was selected and diluted as volumetric ratio within the wells. Testing groups were combined of control samples as non-loaded niosomes and only PTX. Testing samples were prepared at the concentration of 0.0125 mg/mL (the most promising PTX load, see *Supplementary Figure 12*) PTX loaded into niosomes (0.2 mM) and diluted to 1:100. All niosomal dilutions were prepared in the starvation media after 1-day cellular attachment. The number of cells seeded through the wells were 5000 cells/well in 100 μL with overnight incubation at 37°C. 4 serial dilutions from one stock PTX-loaded niosomes (0.2 mM) were exposed to HDF and U87 cell lines for 48 hours. Then, 3-(4,5-Dimethylthiazol-2-yl)-2,5-Diphenyltetrazolium Bromide (MTT) was dissolved in PBS at pH: 7.4 with a final concentration of 0.5 mg/mL, applied 50 μL on each well of the plated, incubated 4 hours. Reading of the wells was performed by removing media and adding 180 μL DMSO (Merck, Germany). The results of cell viability assays were obtained using a microplate reader, detected at the wavelength of 570 nm. (BioRad, USA).

### Flow cytometry assay

U-87, GBM cell lines were seeded in 24-well plates (ISOLAB Co.) at a seeding rate of 2 × 10^4^ cells/well. Seeded cells are adapted overnight before the assay. FITC-tagged niosomes were diluted 1:5000 and added as 1:100 ratio of total volume of the well. Passive incubation was followed by the intervals of 2, 4, 6, 8, and 10 hours and collected separately with their duplicates, demonstrated in *Figure 8*. Trypsin (Pan Biotech) was used to remove the cells for replacing medium by centrifuge at 2000 rpm, 5 mins with PBS and repeating this step continuously three times. Final solution of cell-uptake niosomes are dissolved in PBS containing 1% PFA. After optimization, total cellular uptake was performed after 6 hours of incubation. Cell uptake analysis of each formulation was tested using FACS (BD, USA) with FITC-tagged niosomes. All cell uptake analyzes, including the control group, were analyzed using FlowJo^®^ software and structured by layout mode with histogram-based gating.

### Statistical analysis

*Prism* (License number: GPWF-045319-RIE-6746 to Sabanci University, Version 5.01) was utilized for experimental statistical evaluations. One-way, two-way, three-way ANOVA analysis with one sample t-test confirmations were used for data analysis. Cell viability assay was analyzed by Graph Prism^®^ log-response sigmoidal analysis mode.

## 3. Results & Discussion

### Optimization of niosomal nanoparticle formulations and initial physicochemical characterizations

Although many studies have focused on optimizing the process for drug-loaded niosome formulations in terms of EE% and release assays^(3),(44)–(46)^, few studies have been conducted to understand the physicochemical properties of niosomes with or without drug load.^(12),(16)^ We started with a screening of the production parameters of niosomes and their mutual synergistic effects. We used the thin-layer film hydration method for niosome production, with each step contributing to the variables and constants of physicochemical characterization. These process parameters included surfactant type, constituent molar ratios, number of surfactants, final suspension concentration (mM), sonication time (min), transition temperature (°C), rotation speed (rpm), organic solvent type, final suspension volume (mL), hydration buffer, and hydration volume (mL). However, the effects of many of these process parameters on the physicochemical properties of niosomes, such as size, polydispersity, and zeta potential, were unknown or had not been published before.

To perform an initial screening of the process parameters, we conducted an experiment with niosome production parameters as listed in *Table 1*. We divided the formulations into single-surfactant type formulations (SSTF) and multi-surfactant type formulations (MSTF), each with four different concentrations (mM) and two different sonication times (min). The temperature was 60°C, the rotation speed for the hydration step was 150 rpm, the organic solvent was chloroform, hydration buffer was phosphate buffer saline (PBS), and the hydration volume was 10 mL throughout the study. Our main objective was to understand the effects of concentration (mM), sonication time (min), and surfactant types (MSTF vs. SSTF) on the physicochemical properties of niosomes such as size (nm), PDI, and zeta potential (mV). *Figure 3* shows the overall size (nm) distribution as a function of all process variables across SSTF and MSTF of niosomes, while *Supplementary Figure 2* indicates size distribution across five different formulations at two different sonication times (45 mins vs. 90 mins). *Supplementary Figure 1* shows the polydispersity and zeta potential (mV) distribution profiles at different concentrations (0.1 mM, 0.2 mM, 0.3 mM, and 0.42 mM) for all formulations. To consider concentration variable (controlled) to size variable (uncontrolled), two-way ANOVA statistical consideration is applied. By that, the effect of concentration on size was aimed to be determined. To consider the separate investigation for sonication time on the concentration variable, while sonication time was already having two different set points, unpaired t-test was applied between size and sonication time to investigate the correlation between. The effect of sonication time on the size of five different formulations is shown in *Supplementary Table 1. Supplementary Table 3A* and *3B* provide further details on two-way ANOVA for the PDI and zeta potential of the preliminary formulation screening of niosomes, in addition to sonication time versus size consideration.

**Table 3.** DoE optimized formulation, ingredient, and condition list.

| **Molar Ratio (%)**<br>**(CHO: S60: T80)** | **Predicted Size (nm)** | **Predicted Mixture**<br>**Concentration (mM)** | **Predicted Sonication**<br>**time (min)** | **Obtained Size (nm)** |
| --- | --- | --- | --- | --- |
| **34.5: 43.5: 22** | 100.6 | 0.228 | 63 | 127.7 ± 7.11 |
| **41: 2: 57** | 99 | 0.126 | 47 | 125.4 ± 2.13 |
| **40: 3: 57** | 92.5 | 0.1 | 45 | 114.7 ± 14.14 |
| **49: 49: 2** | 97 | 0.125 | 86 | 142.9 ± 11.32 |
| **49: 49: 2** | 92 | 0.1 | 83 | 149.3 ± 12.11 |
| **48: 4: 48** | 99.8 | 0.11 | 50 | 110.3 ± 9.22 |
| **49: 48: 3** | 99.5 | 0.13 | 89 | 176.7 ± 11.12 |
| **50: 50: 0** | 84.4 | 0.1 | 90 | 141.3 ± 15.48 |
| **50: 50: 0** | 92 | 0.1 | 45 | 137.3 ± 9.23 |
| **50: 50: 0** | 97 | 0.2 | 90 | 145.6 ± 5.34 |

**Figure 3.**
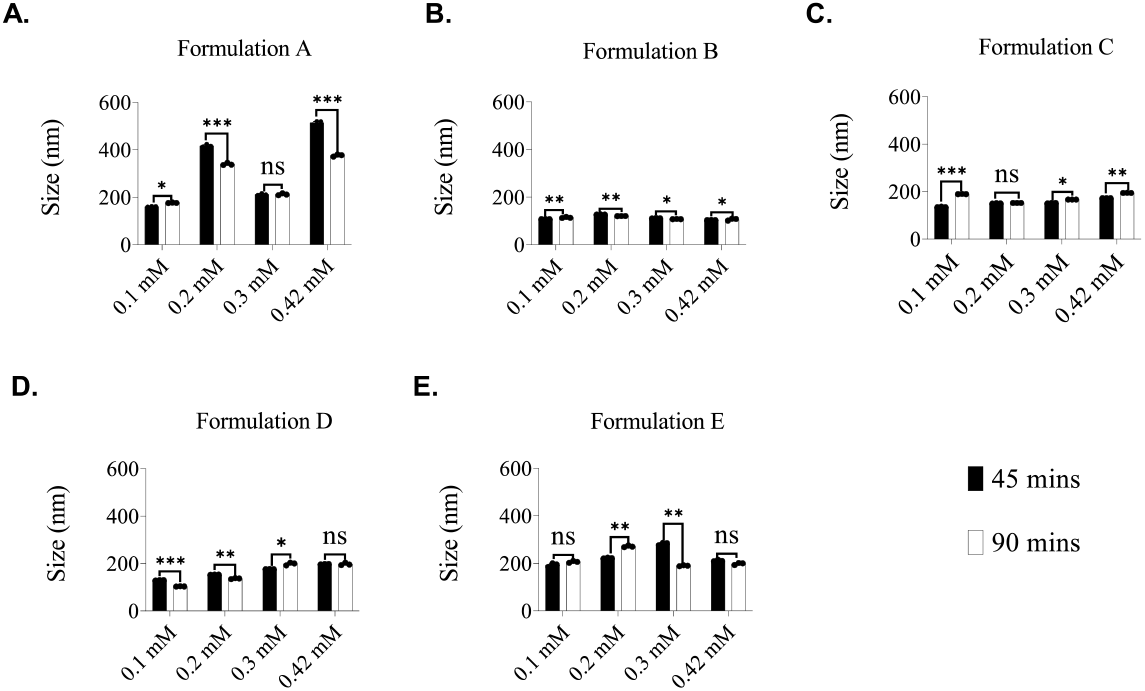
Preliminary effect analysis of sonication, final mixture concentration, and surfactant type profiled for all formulations (A-E), demonstrated in *Table 1*. Two-way ANOVA analysis is applied to all preliminary formulations to identify concentration (mM) and sonication time (min) on size (nm). ns: p-value > 0.05; *: p-value ≤ 0.05; **: p-value ≤ 0.01; ***: p-value ≤ 0.001, ****: p-value ≤ 0.0001

The correlation matrix shown in *Table 2* combines two-way ANOVA and multiple t-tests (*Supplementary Table 2*). The optimal process conditions for niosome production were determined by screening three parameters: 1) Ingredient percentage (%) and type of surfactant, 2) Final mixture concentration, and surfactant type, 3) Sonication time (min). This preliminary screening of niosome production parameters in *Figure 3* guided to determine promising formulation recipes can be statistically optimized via MODDE®. By that, factor-effect analysis was performed. Two-factorial analysis indicated in *Table 2* was formed based on preliminary effect findings in *Figure 3* and on required preset physicochemical specifications. In this evaluation, size, homogeneity and zeta potential were model responses of determined process parameters. According to the preliminary determination, the most significant effect on physicochemical targets of niosome formulations appeared when multiple surfactants were used. Such observation can be observed between Span 60 and Tween 80 correlation. (****: p-value ≤ 0.0001) In addition, Tween 80 has more impact on physicochemical properties than Span 60 correlated with cholesterol, ***: p-value ≤ 0.001 and *: p-value ≤ 0.05, respectively. Although there is no specific effect determined for sonication time or mixture concentration on overall responses, formulation-specific investigation may vary according to obtained results. However, preliminary parameter screening was designed for initial effect evaluation and thus, formulation-specific impacts were not reported. Besides, formulations from A to E were not included to further studies and only used for obtaining statically proposed formulations by MODDE®.

The data obtained were statistically analyzed and further formulations were determined using the DoE program MODDE^®^. The optimized conditions were evaluated by a logarithmic order with the evaluation of the target size between 100–200 nm to ensure the nanoparticles entry into cellular and biological barriers.^(47),(48)^ PDI and zeta potential (mV) were eliminated for the targeting step. MODDE^®^ proposed 10 different optimized experimental conditions prepared based on preliminary findings demonstrated in *Figure 3*. Obtained results of 10 optimized formulations were predicted by MODDE® software based on previously reported data. Predicted and utilized conditions of MODDE® (experimental) were listed in *Table 3*. By applying the proposed conditions by MODDE®, we obtained niosomal nanoparticles, and their size distributions (nm) represented in a single column in *Table 3* and their detailed size (nm), PDI, and zeta potential were shown in *Figure 4*. According to the obtained average size (nm), the overall size range of 10 optimized formulations was between 120 and 180 nm, the PDI range was between 0.1 and 0.3, and zeta potential (mV) range was between -0.5 mV and 0.5 mV. The F-score similarity test was determined as 0.83. Although targeted neutralized surface charge was achieved, stability tracking was required due to their close tendency to be aggregated in such surface properties.

**Figure 4.**
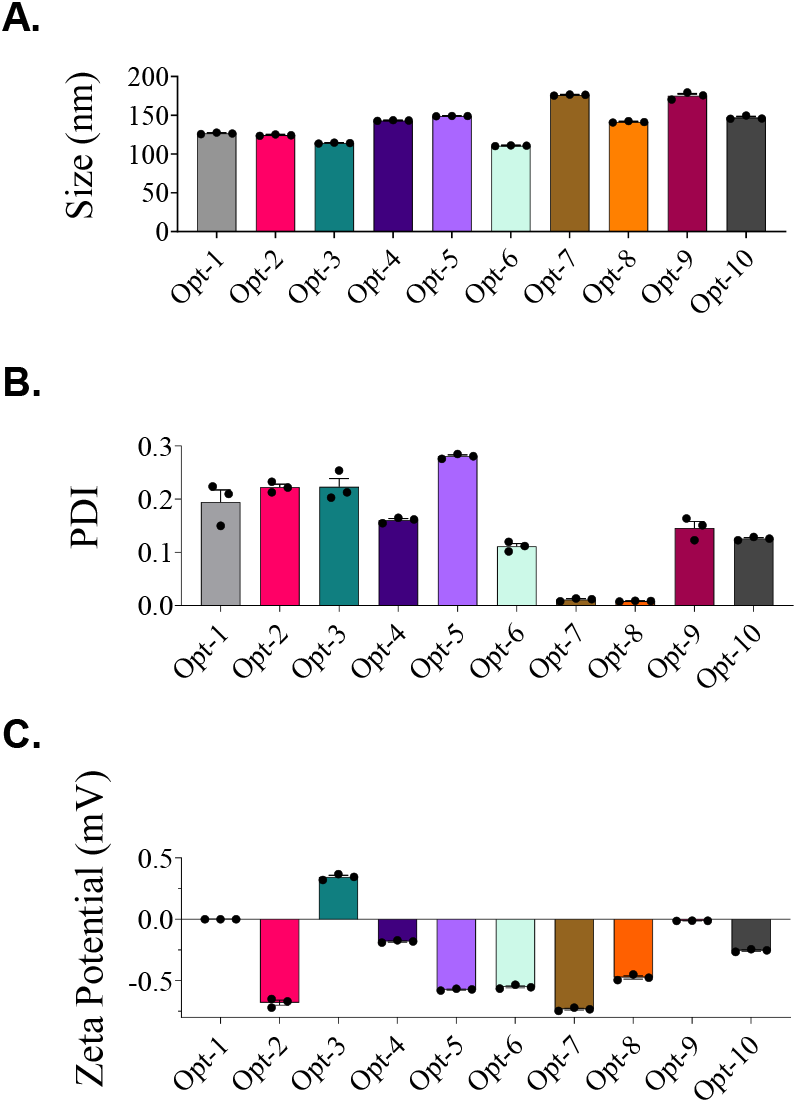
10 different optimized niosome formulation physicochemical profiles A. Size distribution (nm) (Ranging from 100 nm to 200 nm) B. Polydispersity (PDI) (Ranging from 0.05 to 0.3) C. Zeta Potential (mV). (Ranging from -0.5 mV to 0.5 mV) The size distribution of the data is demonstrated in *Supplementary Figure 9*.

### Testing the stability of niosomal nanoparticles

10 different optimized niosome formulations were characterized and found to be in the critical range of size (nm), PDI, and zeta potential (mV). Niosomes are considered to be more stable drug carriers than liposomes, but accurate stability profiles of niosomes have not been established beyond 4 weeks.^(12)^ On the other hand, although size (nm) was followed up to 4 weeks, their PDI and zeta potential (mV) fluctuations by time were not reported. Therefore, a 92-day stability test was performed while storing the samples at +4°C. Each formulation and the variations in its size (nm), PDI, and zeta potential (mV) were compared by applying one-way ANOVA. *Figure 5* shows the change in size (nm) in 92 days for each formulation. Formulations were named according to *Table 3* as the order of columns from Opt-1 to Opt-10. *Supplementary Figure 2* demonstrates PDI and zeta potential (mV) profiles in 92 days for each optimization. The most stable niosome formulations in terms of size (nm) were defined based on slight changes and no aggregation profiles in their peak iterations. The most stable PDI and zeta potential (mV) profiles of niosome formulations were determined based on sudden jumps in PDI and zeta potential (mV). Slight changes were also acceptable decision criteria for PDI and zeta potential (mV). Detailed statistical analysis is presented in *Supplementary Table 4A* and *4B*.

**Table 4:** Drug excapsulation values of PTX-loaded niosomes.

| # | Concentration (mg/mL) | EE% | DLC% | DLE% |
| --- | --- | --- | --- | --- |
| 1 | 0.0125 | 90.3 ± 2.6 | 9.2 ± 2.5 | 89.4± 3.7 |
| 2 | 0.1 | 14.7 ± 3.5 | 0.55 ± 0.06 | 13.2± 1.5 |
| 3 | 0.5 | 9.6 ± 2.1 | 0.11 ± 0.02 | 9.5± 0.3 |

**Figure 5.**
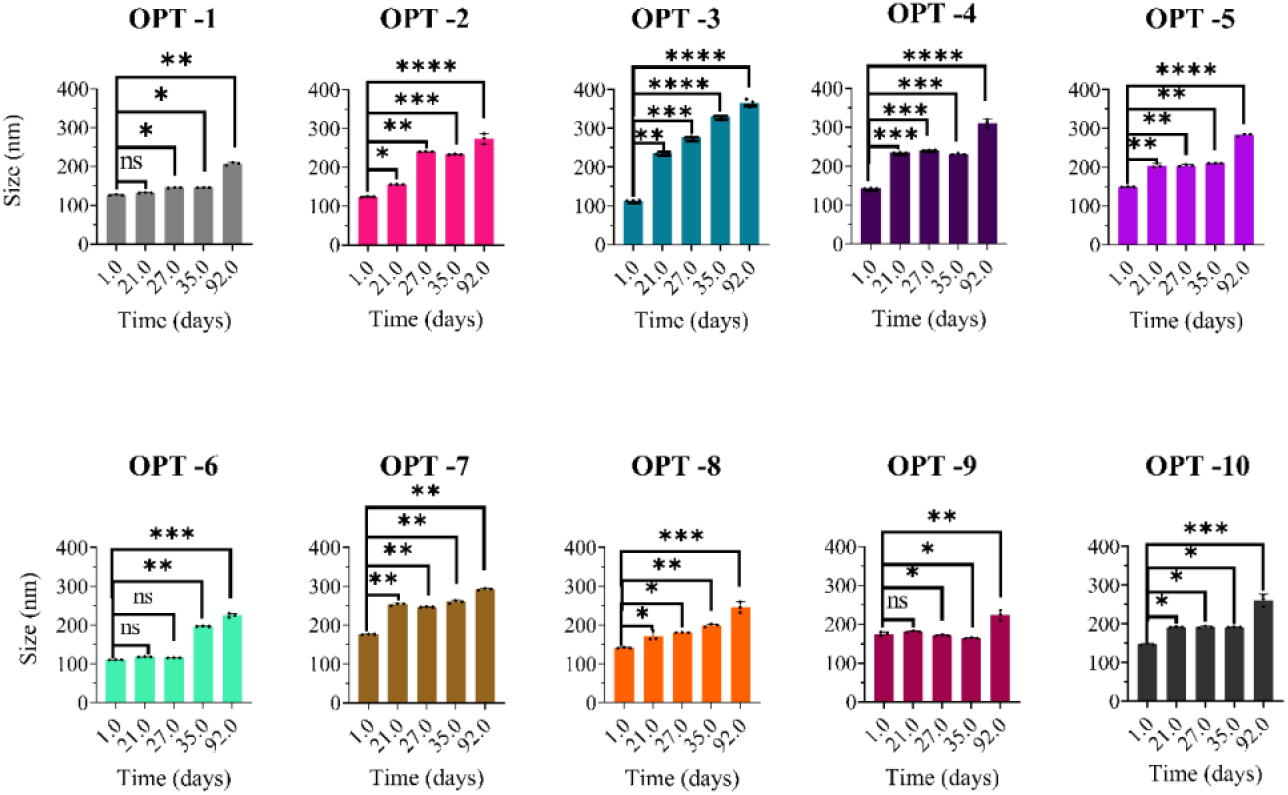
92 days shelf-life profiling of optimized 10 formulations. One-way ANOVA analysis is applied to all niosome formulations to determine overall 92 days shifting profile for A. Polydispersity (PDI), B. Zeta Potential (mV). One sample t-test is applied to each formulation and analyzed for the shift between 1-21 days, 1-27 days, 1-35 days, and 1-92 days in *Supplementary Table 4A* and *4B*. (ns: p-value > 0.05; *: p-value ≤ 0.05; **: p-value ≤ 0.01; ***: p-value ≤ 0.001, ****: p-value ≤ 0.0001.)

*Supplementary Table 4A* and *4B* demonstrated the detailed statistical significance evaluation in their stability profiles for PDI and zeta potential (mV). In the size stability profiles, Opt-1, Opt-6, Opt-8, Opt-9, and Opt-10 showed slight changes in 92 days and were also in the critical size range, which was set at 100–200 (nm). The PDI stability profiles demonstrated that Opt-4, Opt-5, Opt-9, and Opt-10 had slight changes in 92 days. Except for Opt-1, Opt-2, and Opt-3, the PDI of all optimized niosome formulations was in the critical range of 0.2-0.5. The zeta potential (mV) stability profiles were in the critical range of -10 to 10 mV for all niosome formulations. The most stable zeta potential (mV) profiles were selected as Opt-5, Opt-6, Opt-9, and Opt-10 since no fluctuations were observed in their zeta potential measurements within 92 days. Interestingly, Opt-9 was found to have a gradual increase in electronegativity over 92 days among all other optimization formulations, which was still in the range of -10 to 10 mV. Stability tests showed that five niosome formulations were in the critical size range (nm), almost seven niosome formulations were in the critical PDI range, and all niosome formulations were in the critical zeta potential (mV) range in 92 days.

### Testing the reproducibility of niosomal nanoparticles

To determine the reproducibility of the niosome formulations, three experiments were performed at three different days under the same conditions as in *Table 3*. Comparison was made using F-score (Sorensen-Dice coefficient). The F-score test indicates the similarity of each run with respect to the different intervals between days and gave a value of 0.62. The most reproducible data among 10 formulations were obtained with Opt-10, one of the most stable optimized conditions in the size range from 100 nm to 200 nm. The reproducibility test was necessary to use this nanocarrier formulations in drug delivery with greater accuracy. Additionally, the reproducibility test, followed by a 92-day stability test, brought the novel results to the literature where no record of them was found before. *Figure 6* demonstrated the reproducibility profile for the size (nm) and *Supplementary Figure 3* demonstrated the reproducibility profile for the PDI and zeta potential (mV). Based on the reproducibility profiles for the size (nm) of 10 niosome formulations, all formulations except Opt-4, Opt-5 and Opt-7 were found reproducible. Although significant variations were observed in some formulations, they were accepted if the results were found non-excessive within the critical ranges. Detail statistical evaluations can be found in *Supplementary Table 5*.

**Figure 6.**
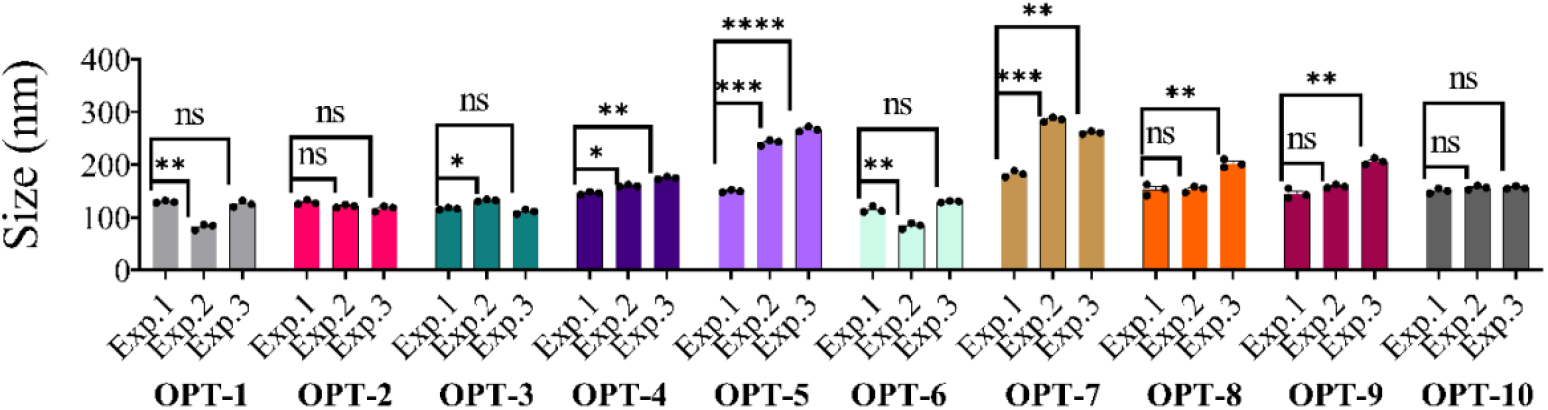
Intraday experiments of 10 optimized formulations demonstrating reproducibility performance of each in size (nm). PDI and Zeta Potential profiles are demonstrated in *Supplementary Figure 3*. Unpaired t-test ANOVA statistical analysis is performed to compare and determine three inter-day experiment group in same experiment conditions. ns: p-value > 0.05; *: p-value ≤ 0.05; **: p-value ≤ 0.01; ***: p-value ≤ 0.001, ****: p-value ≤ 0.0001.

### FTIR profiling for lyophilized niosomes and liposomes

FTIR analyses were performed to select certain optimized and promising niosomal formulations based on corresponded sole ingredient spectra. In *Supplementary Figure 8*, niosomal nanoparticles and liposomes have common chemical groups with several differences. First, all niosomal formulations have demonstrated the similar wavelength ranges from 3600 to 3200 cm^-1^ which represents parallel findings to Span 60 sole spectra demonstrating O-H stretching.^(49)^Another spectral observation is made in the wavelength range of 2400 to 2600 cm^-1^ typically corresponds to the region associated with aliphatic C-H stretching.^(50)^ This range often includes vibrations from methyl (CH_3_) and methylene (CH_2_) groups present in organic molecules. The peaks in this range can indicate the presence and nature of these aliphatic hydrocarbon chains.

L-α-lecithin should give several wavelengths observed in 2800-3000 cm^-1^ which corresponds to C-H stretching vibrations in the hydrocarbon chains. Another wavelength observation is demonstrated ester carbonyl stretch of the phospholipid head group within the wavelength of 1740-1750 cm^-1^. Other phosphate group related vibrations are observed within the wavelength of 1200-1300 cm^-1^. Although niosome and liposome chemical groups indicate the expected wavelength ranges, determined O-H stretching and C-H stretching characteristics are changed by the formulation conditions. For instance, Opt-5 demonstrates the least concentrated O-H stretching compared to other formulations. Besides, in this region, it is clearly seen that O-H stretching is the most affected chemical group by the formulation conditions. This observation suggested the importance of process conditions applied during the synthesis, such as temperature, pressure, or duration of reactions, can affect the structure and arrangement of molecules on the nanoparticle surface, potentially impacting O-H stretching vibrations.^(2)^ On the other hand, Opt-5 is found one of the most affected formulations from shelf-life that refers the more stable and concentrated O-H stretching promotes the stability of niosomal nanoparticles. The O-H stretching vibrations are often associated with surface-bound hydroxyl groups.^(50)^ A decrease in O-H stretching intensity might imply fewer available hydroxyl groups on the surface, which could indicate modifications or changes in the surface chemistry. This alteration might impact the stability of nanoparticles, especially if these hydroxyl groups are involved in stabilizing the nanoparticle dispersion or in interactions with surrounding molecules. While changes in O-H stretching intensity in the FTIR spectrum can provide indications about the stability of nanoparticles, it’s essential to consider multiple factors and conduct complementary analyses to draw comprehensive conclusions about nanoparticle stability and behavior. Thus, such complimentary analysis done through the study provide a deep understanding and robust formulation selection for further therapeutic functionalization.

### Assessing the cellular toxicity of niosomal nanoparticles

Physicochemical characterization of niosome formulations, stability profiles and reproducibility tests were performed, and promising formulations were recorded. To test their cellular toxicity, MTT assay was performed two different cell sources: U-87 and NFS-60. *Supplementary Figure 4* shows the cellular viability profiles of 10 different niosome formulations and liposomes of the glioblastoma cancer cell line U-87. We screened the cellular viability profiles of these niosomal formulations after 48 days of their synthesis to determine whether the range of cellular viability had gradually shifted, as shown in *Figure 7* for U-87. In addition, the leukemia cell line, NFS-60 was tested with the same niosome formulations and liposomes (see *Supplementary Figure 5)*. Both of these different cell lines with the same niosome and liposome formulations showed no cellular toxicity and had similar cellular viability profiles. Cellular viability ranged from 80% to 100% with all ten different niosomal formulations and liposomes when they tested only a day after their synthesis. On day 24 and day 48 after their synthesis, cellular viability ranged from 75% to 100%. The results also confirmed the stability test of the niosome formulations, in almost all niosome formulations were stable for at least 2 months. In order to make final decisions for further selection of optimal niosomal formulations, their cellular uptake profiles needed to be investigated.

**Figure 7.**
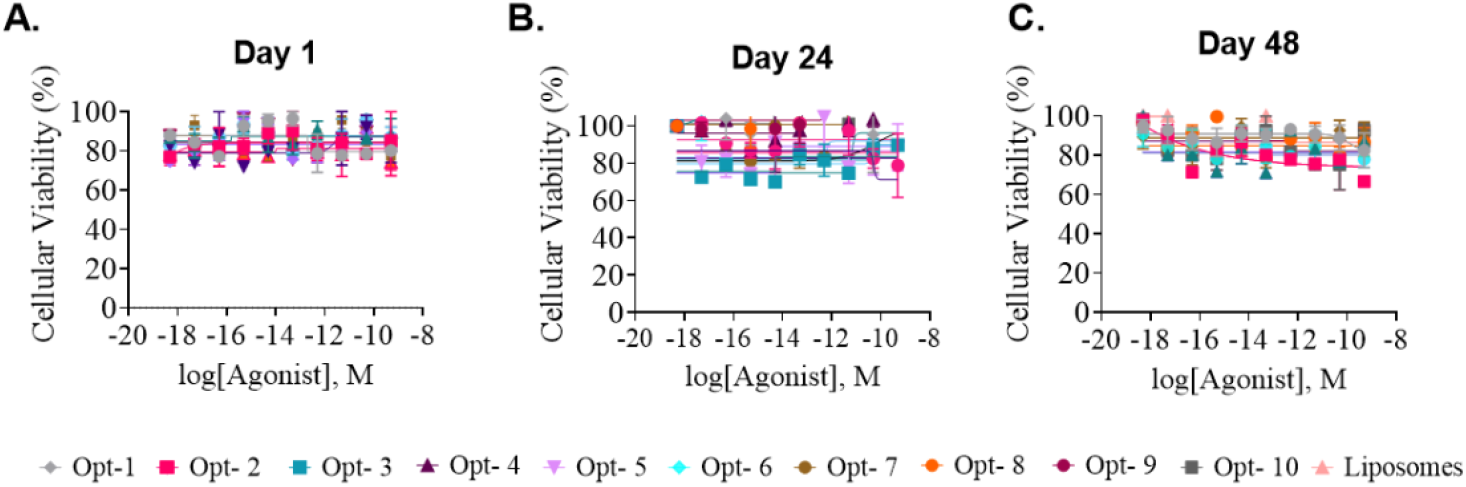
48 days cellular viability MTT assay for all niosome optimizations and liposome comparison performed in U-87 cell line. A. Day 1 cell viability (%) in the range of 80-100. B. Day 24 cell viability (%) slow shift to the range of 75-100. C. Day 48 cell viability (%) slow shift to the range of 75-100. 5000 cells/wells are used, and 9 serial dilutions are applied. Sigmoidal log-response concentration (M) indicates the x-axis.

### Cellular internalization of niosomes

Fluorescently labeled niosomes were generated for cellular assessments. A total of 10 distinct niosome formulations, along with liposomes labeled using the fluorescent tracer FITC, were subjected to analysis through flow cytometry.^(51)^

FITC labelling of niosomes were based on the total volume of niosomal formulations before the production process. The overall, based on the desired last concentration, FITC existence was set to 0.05% of the total volume from the stock solution of 300 nM. The percentage of FITC and stock solution concentration are set, and encapsulation efficiency was calculated based on sole standard curve of FITC. Standard curve of FITC and the linear curve equation was presented in *Supplementary Figure 6*. After encapsulation of FITC, high-speed centrifugal release (12,000 rpm, 20 mins) was used, and supernatant absorbance was recorded triple to calculate the entrapment efficiency. *Supplementary Figure 7* indicates the stability of FITC labelling efficiency between replicates. All encapsulation efficiency results obtained from different FITC concentrations are demonstrated in *Supplementary Table 7*. Results demonstrated almost all optimized niosome formulations are gradually decreased their EE% by increased FITC concentration and not all niosomal formulation demonstrated the same EE% at the same FITC concentration. L-α-lecithin liposomes’ FITC tagging are performed after determining the optimum FITC concentration for niosome formulations.

To utilize the formulated niosomes as improved carrier agents, investigation of their cellular uptake was critical. The cellular source U-87 was used for the study. Incubation time optimization was performed based one of the most reliable physicochemical standings, Opt-10 and the change in FITC-A mean was analyzed by using statistical interface embedded in FlowJo. *Figure 8* is demonstrating the only cell population and corresponded FITC-A mean which could be also considered as baseline point for the overall FITC-A mean intensity. Flow cytometer allows three different runs from the same sample, however intraday uptake profiles cannot be considered as one of the replicates for the same experiment. However, their trend could be comparable. Results demonstrated that after 6-hour incubation, slight decrease in FITC-A mean is observed. Thus, all further cellular uptake profiling of niosome and liposome nanocarriers were tested in 6-hour incubation.

**Figure 8.**
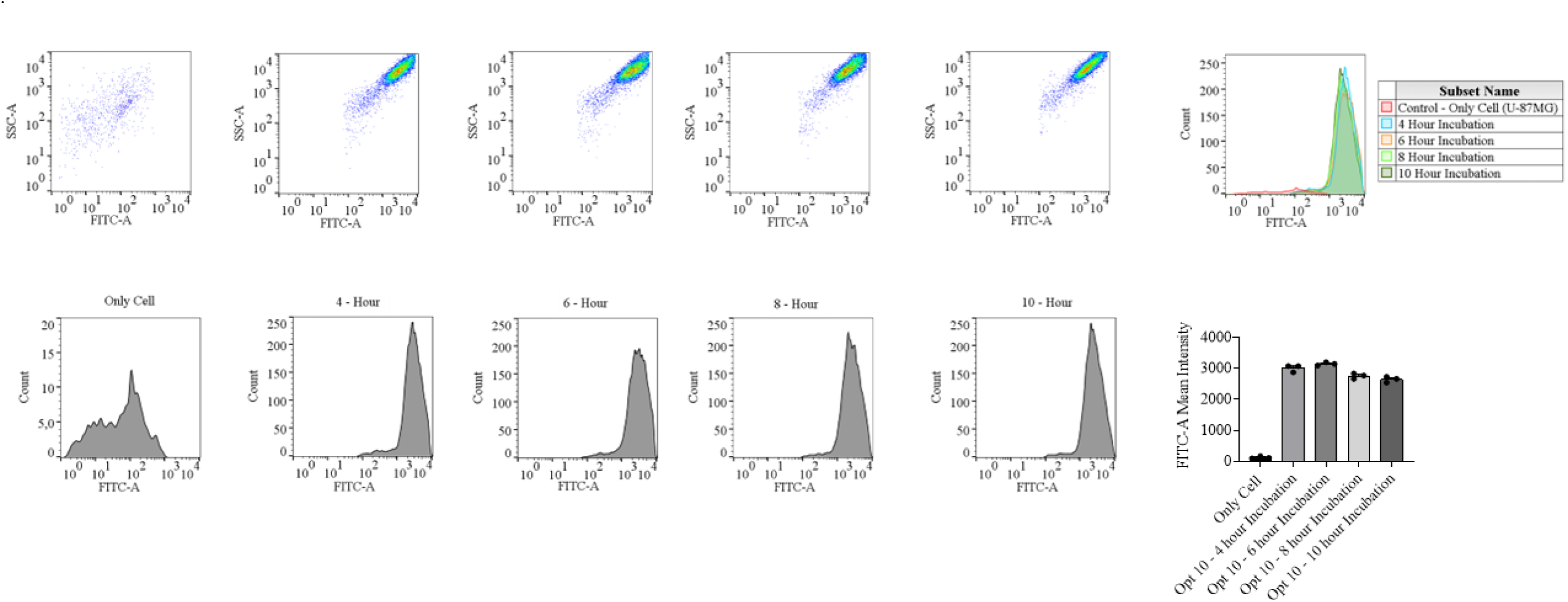
Different incubation time screening profiles for cellular uptake in 4 h, 6h, 8h, and 10 h. FITC-A mean distribution which is normalized based on internal loading control is graphed and selected based on the highest FITC-A mean. Each formulation cellular uptake is obtained as triplicates from the same population, different well-plates. 2 × 10^4^ U-87 cells are used for each FITC-tagged niosomes.

The incubation time optimization demonstrated that long incubation period could diminish the uptake could be caused by decreased intensity of FITC since 48-hour cellular viability indicated no toxicity observed for all niosome formulations for U-87MG and NFS-60 cell lines. After determination of the most proper incubation time, simultaneous cellular uptake profiling for niosomal and liposomal formulations were tested for 6 – hour. *Figure 9* is demonstrating individual FITC-A mean captured for all testing groups.

**Figure 9.**
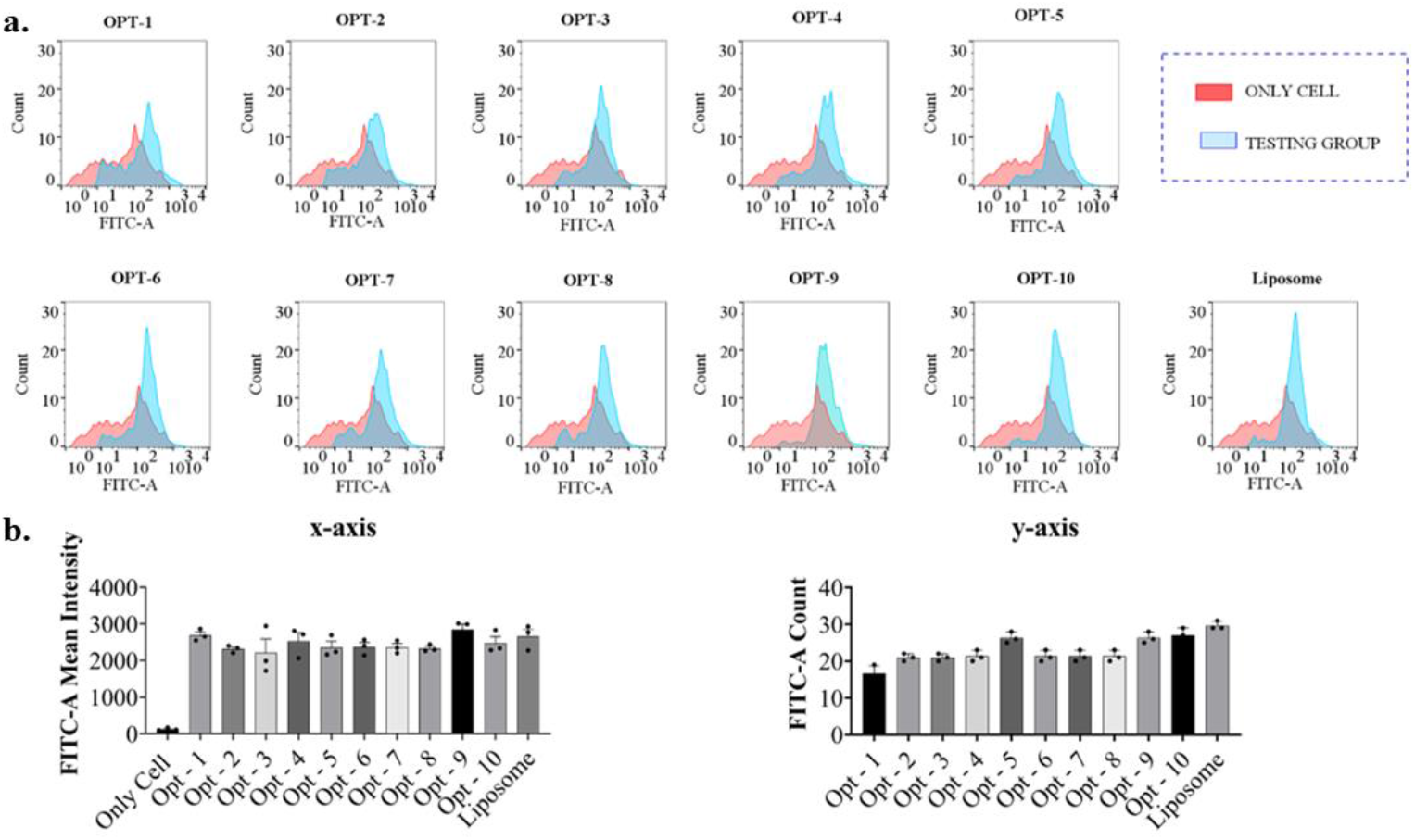
Simultaneous cellular uptake analysis performed for niosome (Opt-1 to Opt-10) and liposome (L-α-lecithin) formulations. Testing groups and a control group were represented as light blue and light red, respectively. **a**. x-axis and y-axis shifts of test samples and their histogram overlay with only cell population. **b**. Normalized FITC-A signal evaluation based on x-axis and y-axis shift.

The analysis conditions were set based on the sole population FITC-A mean intensity which was asset for the starting gate. The sole population was supposed to not have FITC-A count. However, FITC autofluorescence positive feed is already reported in the literature.^(52)–(54)^ Thus, the recorded FITC-A intensity in sole cell population was considered as autofluorescence of FITC abled laser taken as baseline. In the evaluation, not only forward scattered light could be enough for picking the proper formulation. The results demonstrated that x-axis or forward scattered light determined uptake is not indicating any advance or difference, means cells are not granulized and/or swell out during the uptake. If so, they are not demonstrating any difference between each other. On the other hand, site-scattered light versus FITC-A mean indicating that, glioblastoma cells are able to uptake the niosomal nanoparticles in the y-axis manner. It means the uptake should be analyzed in both ways as it is demonstrated in *Figure 9*.

Cellular internalization analysis initiated with seeking for optimum internalization time sets that were not previously reported. To utilize the same internalization time, several simultaneous analytical considerations were made. Initially, same FITC amount was utilized to all nanocarriers to not cause any false positive signal results. The similar consideration is accomplished in seeded cell number equality. Since seeded concentration and incubation time in their internalization were also same, the only variance was nanocarrier formulations itself. In *Figure 9*, x-axis evaluation was normalized and subtracted from the only cell FITC-A signal. However, x-axis change indicates overall granulation inside the cell after uptake of nanocarrier. Thus, the change in x-axis become normal and indicate that all tested niosomal and liposomal nanocarriers were having similar uptake performances that made the decision even harder. On the other hand, there were significant signal differences observed in y-axis and thought to be alternative path to evaluate nanocarrier performance difference between niosomal and liposomal nanocarriers. Evaluation in y-axis could give the idea of attachment for nanocarriers on the surface of cells and could become longer in length of cells. Therefore, y-axis evaluation was normalized and graphed by subtracting only cell FITC-A signal.

Based on all these different analysis for stability, reproducibility, cytotoxicity, and cellular uptake analyses, we found that Opt-6, Opt-8, Opt-9, and Opt-10 were selected as the best formulations among all 10 different formulations (*Supplementary Figure 9)*. The cellular uptake results showed that even though liposomal and niosomal formulations had similar cellular viability profiles. The study also found that these four niosomal formulations were in the appropriate range in physicochemical characterization, stability, and reproducibility studies for further cellular applications.

Obtained results from flow cytometry and internalization assays suggest four niosome formulations indicate similar performance with liposomes. To confirm the obtained results with visual data, confocal scanning fluorescence microscopy (CSFM) is used. FITC-tagged Opt-6, Opt-8, Opt-9, and Opt-10 are observed and compared with liposomes simultaneously. *Supplementary Figure 10* demonstrates recorded data for FITC-tagged nanoparticles and proposed enhanced internalization, especially as observed in Opt-10.

### PTX Loading to Niosome

Following the identification of optimal noisome formulation, Opt-10, we proceeded to load it with the chemotherapeutic agent, PTX, to evaluate its drug loading capacity and drug delivery efficiency. Based on pervious studies, PTX-loaded niosomes are prepared at concentration ranging from 0.5 to 1 mg/mL. ^(56)(57)^ However, other studies have reported initial encapsulation of PTX-loaded niosomes at lower concentrations, such as 10 μg/mL.^(58)^ In this study, we tested the encapsulation efficiency (EE%) of Opt-10 using three PTX concentrations: 0.0125, 0.1, and 0.5 mg/mL. To calculate the EE%, a standard curve of PTX was generated with six different concentrations (*Supplementary Figure 12*.*a*.). The resulting graph equation was applied to determine the amounts of encapsulated and free PTX. Although Opt-10 niosomes maintained their size within the desired range, the EE% decreased significantly as the PTX concentration increased from 0.0125 mg/mL to 0.5 mg/mL (*See Supplementary Figure 12*.*b*.). The physicochemical properties of PTX-loaded niosomes were analyzed using DLS, and the size distribution is depicted in *Figure 10a* and *Supplementary Figure 13*.*a-d. Table 4* presents the EE%, DLC%, and DLE% of PTX-loaded niosomes, calculated using the equations provided in the Methods section (Eq. 1-4).

**Figure 10.**
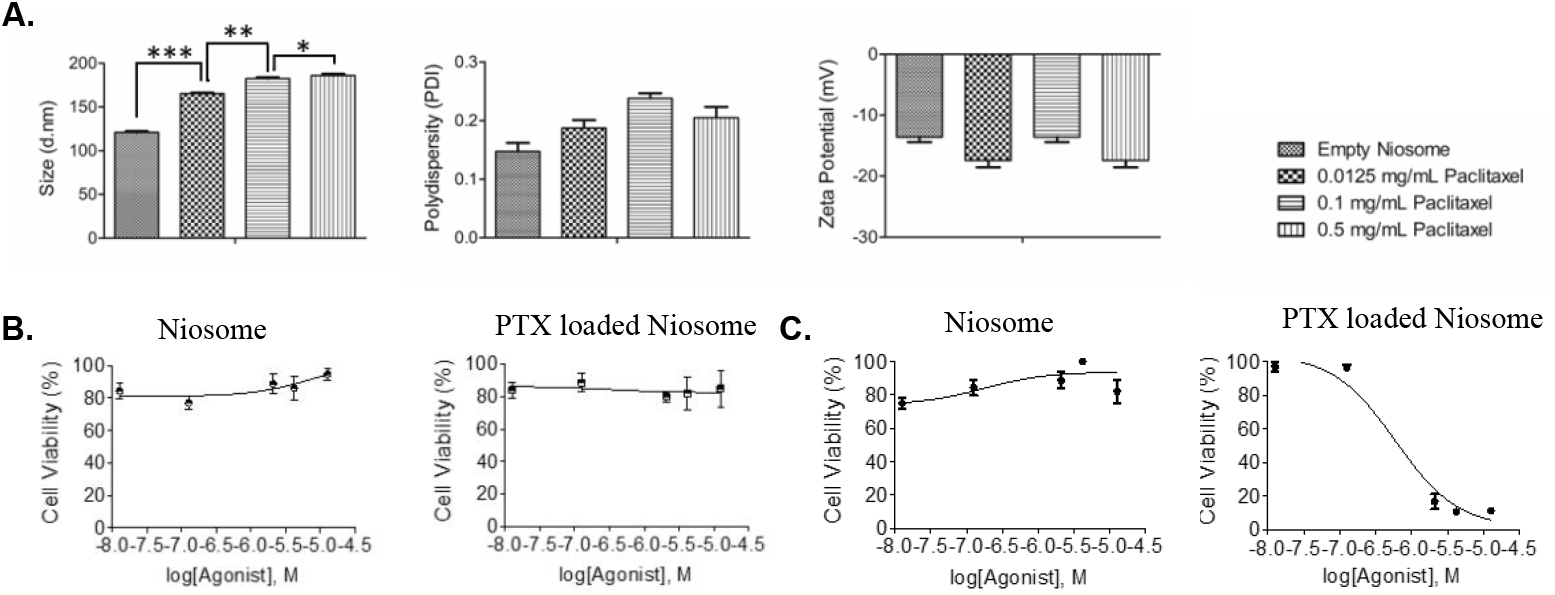
a. DLS and Zeta Potential measurements of empty and PTX loaded noisome (Opt-10). Error bars indicate Standard deviation, and statistical test were performed by ANOVA with Tukey’s multiple comparison; * P<0.05, **P<0.001, *** P<0.0001. b. Cell viability of empty noisome and PTX-loaded noisome (Opt-10) on HDF. c. Cell viability of empty noisome and PTX-loaded noisome (Opt-10) on U87.

The results indicate that Opt-10 niosomes cannot be loaded with more than 0.5 mg/mL PTX without exceeding the target size range of 100 to 200 nm (*Figure 10*). The lowest concentration (0.0125 mg/mL) was choosen for subsequent experiment, as it achieved the highest EE% while maintaing the target size range (121 ± 12 nm) under optimized condition. Furthermore, we compared the cytotoxicity of unloaded noisomes and PTX-loaded noisomes on human-derived fibroblasts (HDFs) (*Figure 10b*) and U87 (Figure 10c). The results showed that unloaded niosomes exhibited no toxicity toward either cell line, while PTX-loaded noisomes were selectively cytotoxic to U87 cells, without affecting the healthy fibroblasts.

## 4. Conclusion

This study rigorously optimized niosomal nanoparticle formulations, conducting in-depth analyses of their physicochemical properties and cellular interactions, notably addressing a crucial research void through a comprehensive comparative examination alongside liposomes. Exploring diverse process parameters, it delineated pivotal size, polydispersity, and zeta potential ranges crucial for optimal niosome design, leveraging MODDE® software to derive ten refined formulations.

Significant findings highlighted the pronounced influence of surfactant types and the synergistic impact of concentration and sonication time on formulation outcomes. Importantly, this research identified indispensable size (100–200 nm), polydispersity (0.2–0.5), and zeta potential (−10 to 10 mV) thresholds essential for superior niosomal performance, bridging an important gap in simultaneous liposome-niosome comparisons. Extended stability assessments spanning 92 days demonstrated promising shelf-life profiles, showcasing formulations adhering consistently within specified critical ranges. Cellular studies revealed minimal toxicity and efficient uptake in select optimized niosomal formulations (Opt-6, Opt-8, Opt-9, and Opt-10), comparable to liposomes.

These refined niosome formulations, boasting stability, reproducibility, low toxicity, and efficient cellular uptake akin to liposomes, signify their potential as economical and versatile nanocarriers. This pioneering comparative investigation with liposomes fills a significant research void, potentially reshaping drug delivery strategies and substantially elevating treatment efficacy and safety.

## Supplementary Information

Supplementary information is available online, which includes the following data and profiles for niosomal formulations: Polydispersity and zeta potential profiles of niosomal formulations; Size distributions of different niosome concentrations; Polydispersity and zeta potential profiles of interday experiments; Cellular viability of U87 cells and NFS-60 cells; Standard curves for FITC labeling of liposomes and niosomes; Flow cytometry data for FITC-labeled niosomes; FTIR analysis of liposome and niosome formulations; Schematic summary for niosome formulation characterization; Confocal images; Flow cytometry data at different incubation times of niosomes.

Tables listing detailed data for the analysis of the effect of sonication time by increased concentration for five different formulations; Correlation analysis of formulation parameters for niosome production; Encapsulation efficiency and FITC concentration optimizations; Statistical analysis results for concentration and sonication time parameters based on pre-set criteria of size, PDI, and zeta potential; One-way ANOVA test results for the effect of the concentration parameter on PDI and zeta potential for each formulation; Stability profiles of optimized formulations in terms of PDI for 21 days, 27 days, 35 days, and 92 days; Stability profiles of optimized formulations in terms of zeta potential for 21 days, 27 days, 35 days, and 92 days; Reproducibility profiles of optimized formulations in size (nm), PDI, and zeta potential (mV); Dynamic Light Scattering data for liposomes.

## Acknowledgement

Authors would like to express their gratitude with the Faculty of Engineering and Natural Sciences (FENS) at Sabanci University for their support, Marie Curie Action Widening Fellowship for funding on the project number 101028391 (to N.M), EMBO Installation Grant for funding on the project number IG-5352-2023 (to N.M.), TUBITAK 2244 Industrial PhD Fellowship under the project number 118C149 (to N.C). This work was also supported by the BAGEP Award of the Science Academy (to N.M.). Many thanks to ILKOGEN Biotech Company for providing instruments and facility.

A preprint of the work has previously been reposited online^59^.

## Data Availability

All data generated or analyzed during this study are included in this published article (and its supplementary information files), or are available from the corresponding author upon request.

