## Supplementary Info for "Optimizing Niosome Formulations for Enhanced Cellular Applications: A Comparative Case Study with L-α-lecithin Liposomes"

**Supplementary Figures**

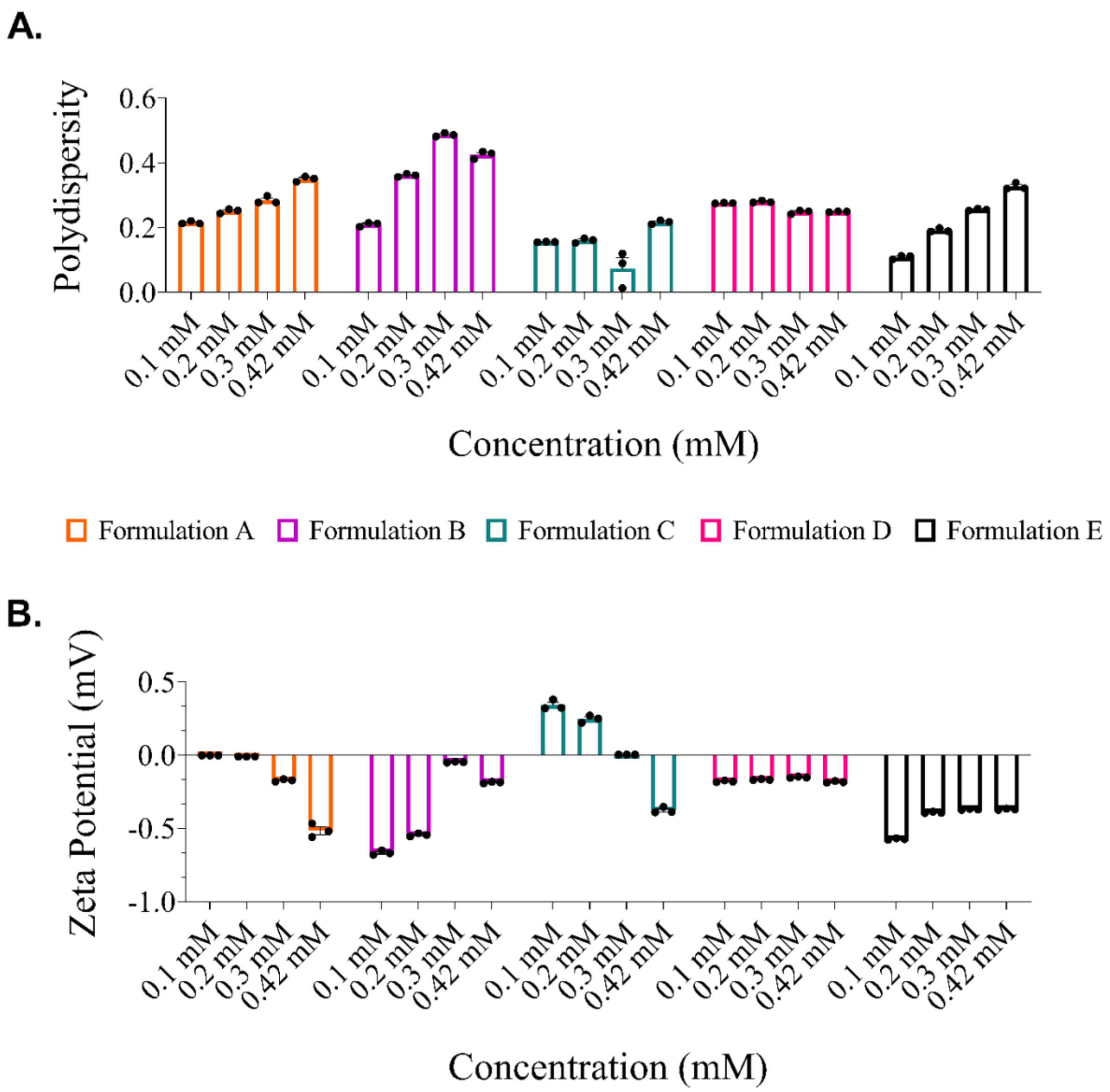


*Supplementary Figure 1: Polydispersity and zeta potential (mV) distribution profiles in different concentrations (0.1mM, 0.2 mM, 0.3 mM, and 0.42 mM) for all formulations. One-way ANOVA is applied to determine the effect of concentration (mM) on size (nm). ns: p-value > 0.05; *: p-value ≤ 0.05; **: p-value ≤ 0.01; ***: p-value ≤ 0.001, ****: p-value ≤ 0.0001. Supplementary Table 4A and 4B demonstrates one-way ANOVA in detail for PDI and Zeta Potential (mV) of niosome preliminary formulation screening*

**A.**

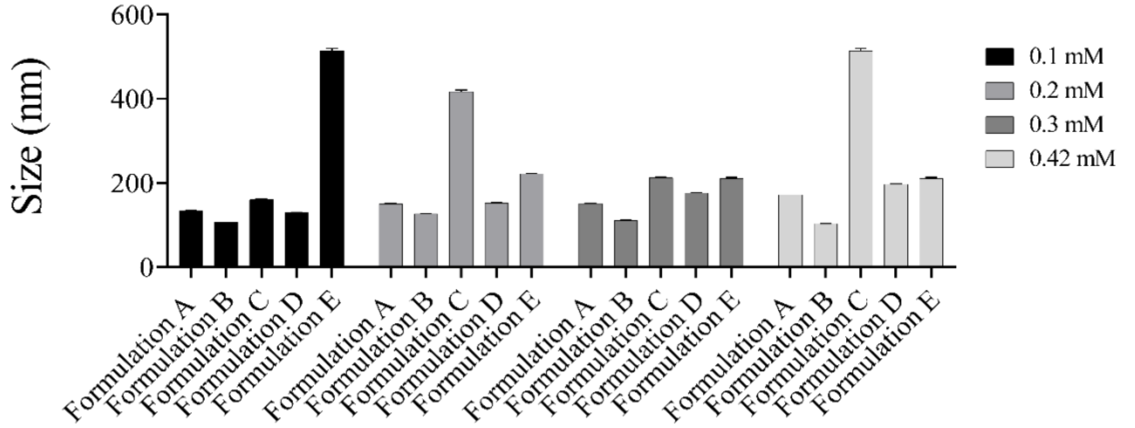


**B.**

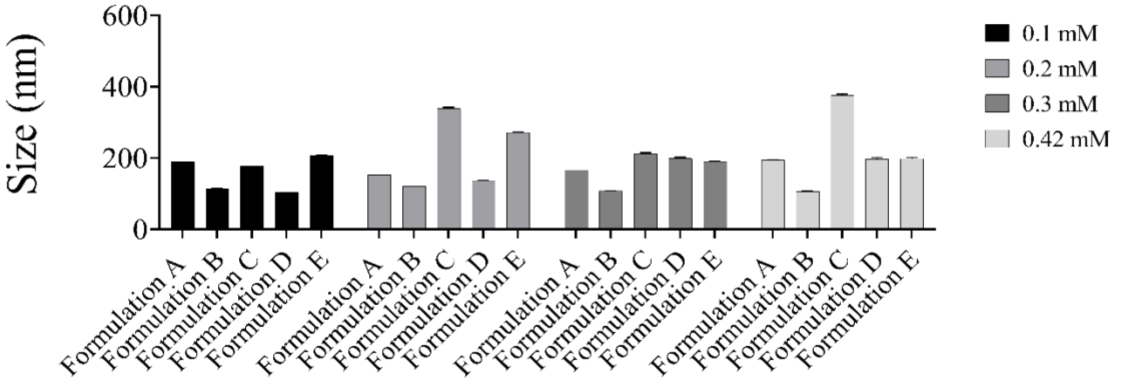


*Supplementary Figure 2: Size (nm) distribution over four different concentrations (0.1 mM, 0.2 mM, 0.3 mM, and 0.42 mM) A. 45 mins, B. 90 mins.*

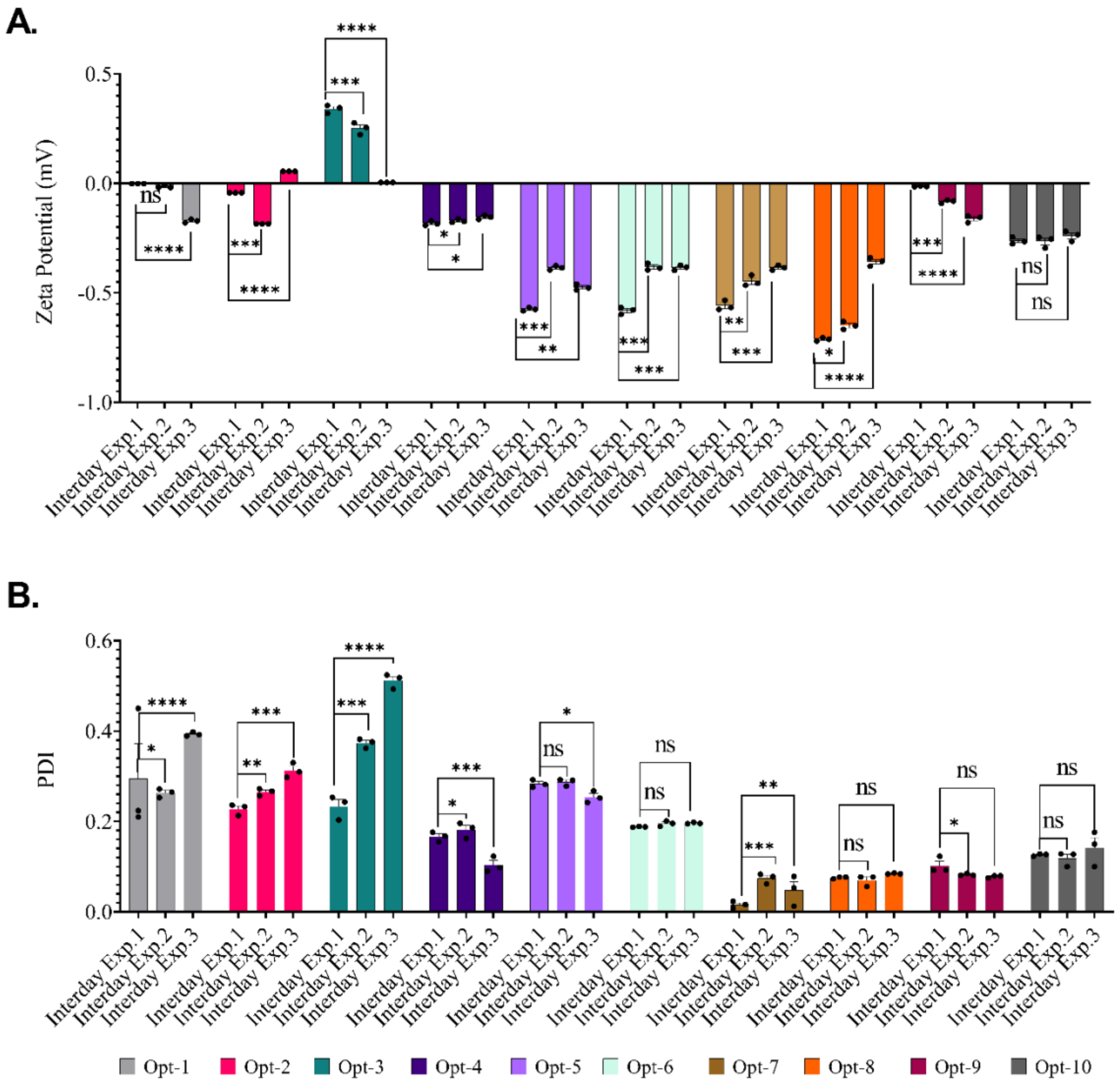


*Supplementary Figure 3: Inter-day experiments of 10 optimized formulations demonstrating reproducibility performance of each in A. Zeta Potential, B. PDI profiles are demonstrated Three -way ANOVA statistical analysis is performed to compare and determine three inter-day experiment group in same experiment conditions. ns: p-value > 0.05; *: p-value ≤ 0.05; **: p-value ≤ 0.01; ***: p-value ≤ 0.001, ****: p-value ≤ 0.0001*

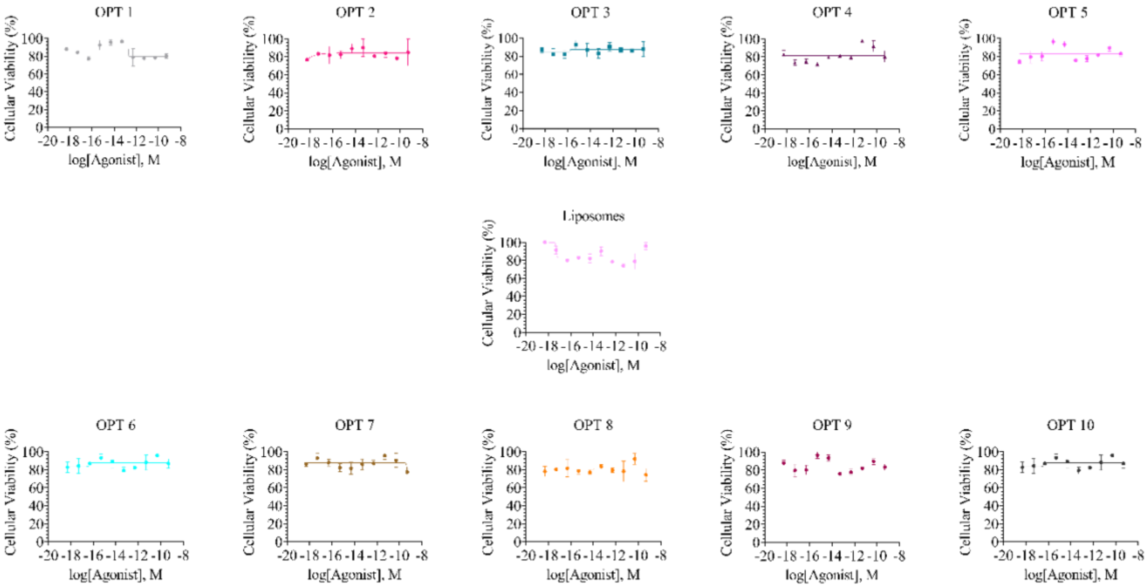


*Supplementary Figure 4: Individual cellular viability MTT assay for all niosome optimizations and liposome comparison performed in U-87 cell line. 5000 cells/wells are used, and 9 serial dilutions are applied. Sigmoidal log-response concentration (M) indicates the x-axis.*

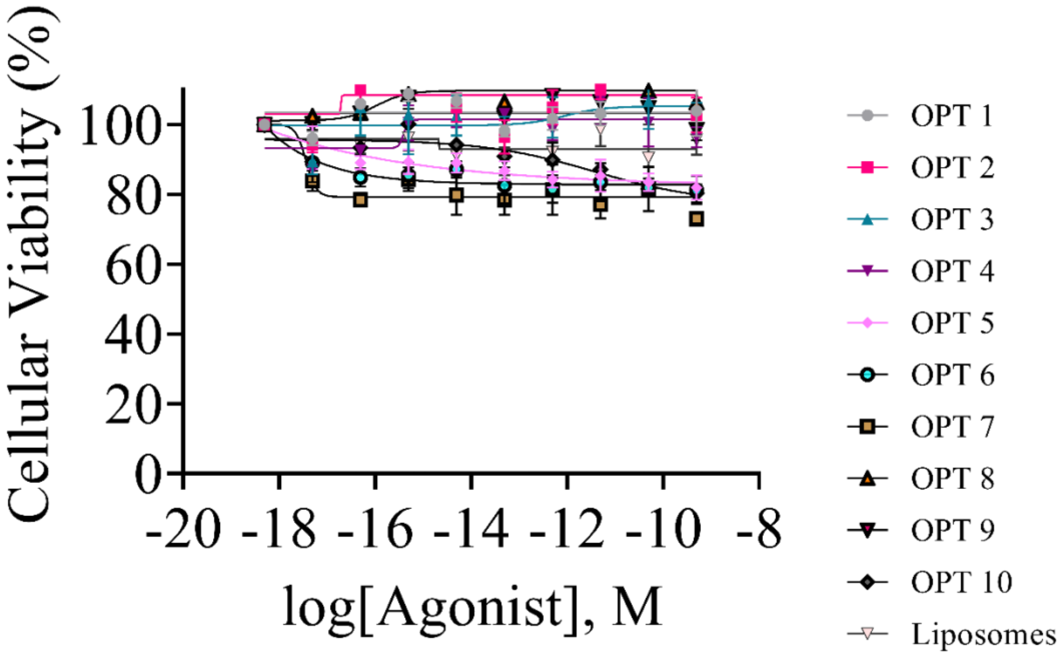


*Supplementary Figure 5: Individual cellular viability MTT assay for all niosome optimizations and liposome comparison performed in NFS-60 cell line. 5000 cells/wells are used, and 9 serial dilutions are applied. Sigmoidal log-response concentration (M) indicates the x-axis.*

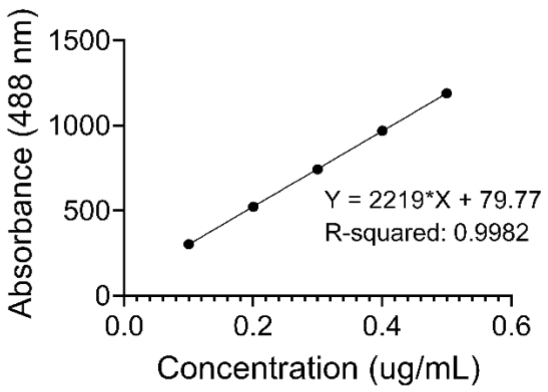


| Concentration (ug/mL) | Absorbance (488 nm) | | |
|---|---|---|---|
| X | A:Y1 | A:Y2 | A:Y3 |
| 0.1 | 300.223 | 305.485 | 303.414 |
| 0.2 | 522.523 | 525.613 | 520.412 |
| 0.3 | 744.823 | 741.165 | 742.810 |
| 0.4 | 967.123 | 970.162 | 969.180 |
| 0.5 | 1189.420 | 1190.250 | 1188.780 |

*Supplementary Figure 6: FITC standard curve to calculate encapsulation efficiency (EE%) of optimized niosomal formulations.*

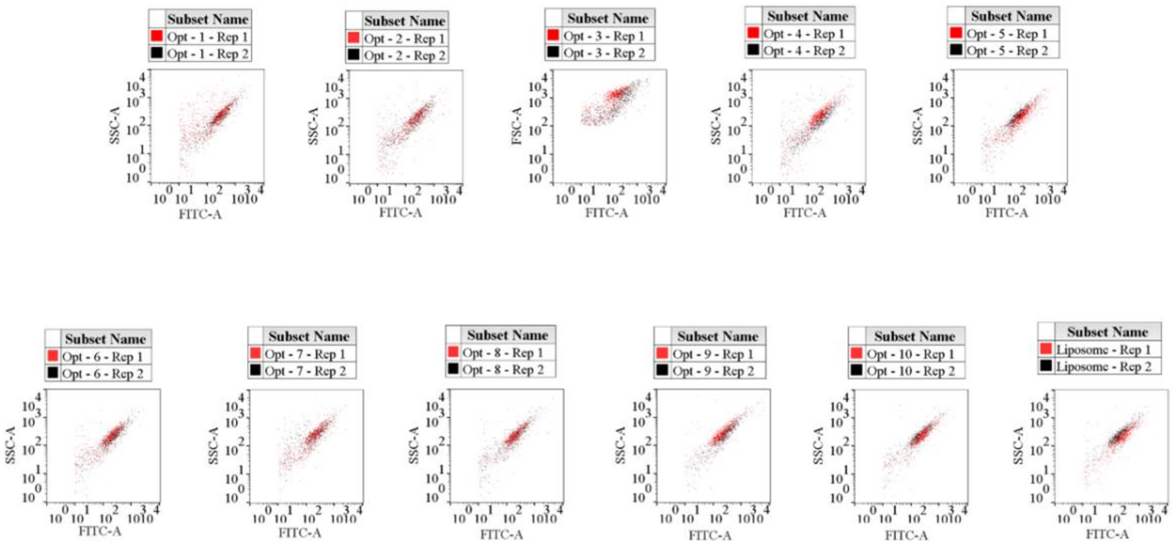


*Supplementary Figure 7: Individual FITC-A mean shifts versus control group for all noisome formulations (Opt-1 to 10) and liposome. Profiles are obtained from FlowJo Software.*

.

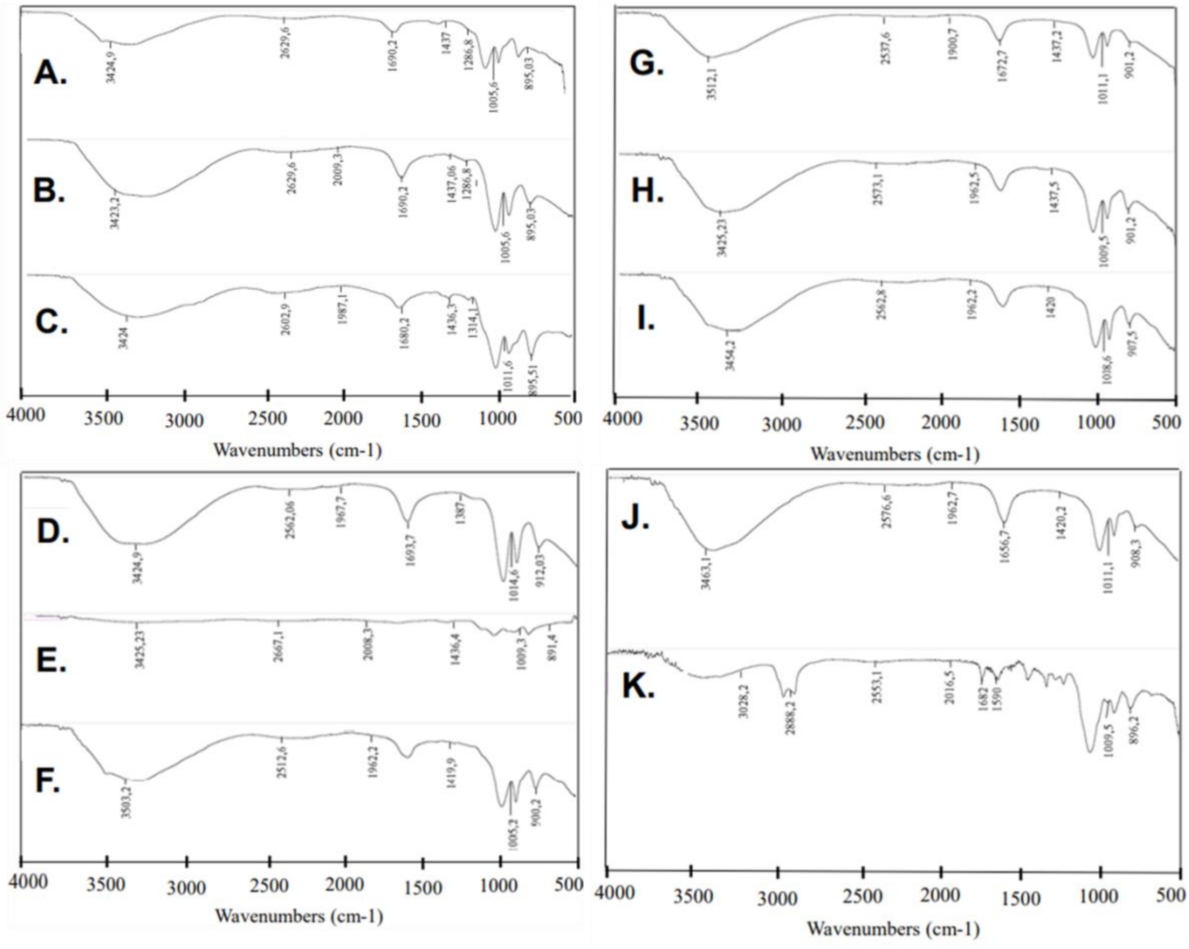


*Supplementary Figure 8: Individual FTIR data of liposome and niosome formulations A. Opt-1, B. Opt-2, C.Opt-3, D. Opt-4, E. Opt-5, F. Opt-6, G. Opt-7, H. Opt-8, I. Opt-9, J. Opt-10, K. Liposome*

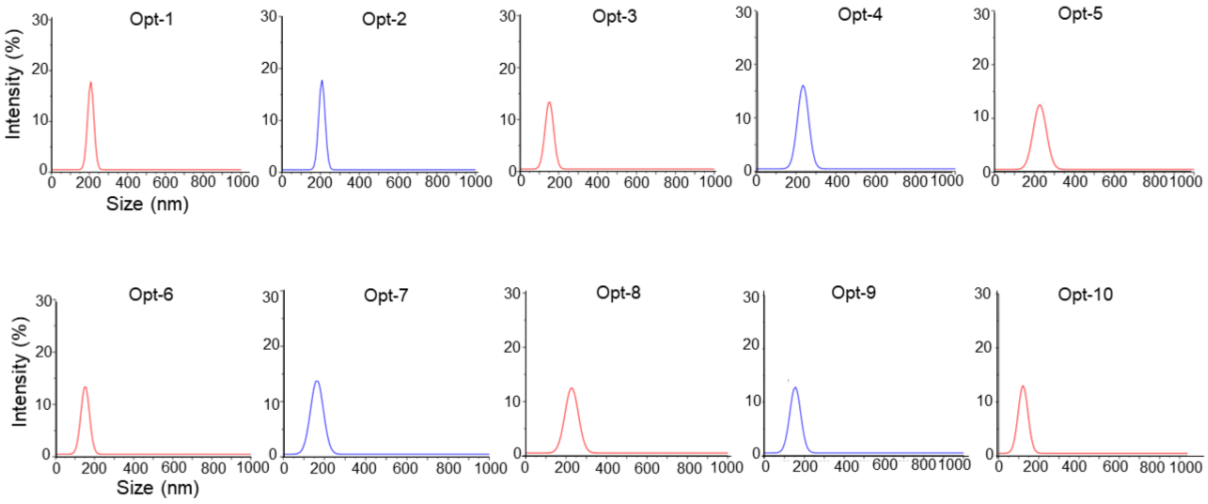


*Supplementary Figure 9: DLS size profiles graphed by Origin® presenting their fresh distribution for each optimized niosomal formulations.*

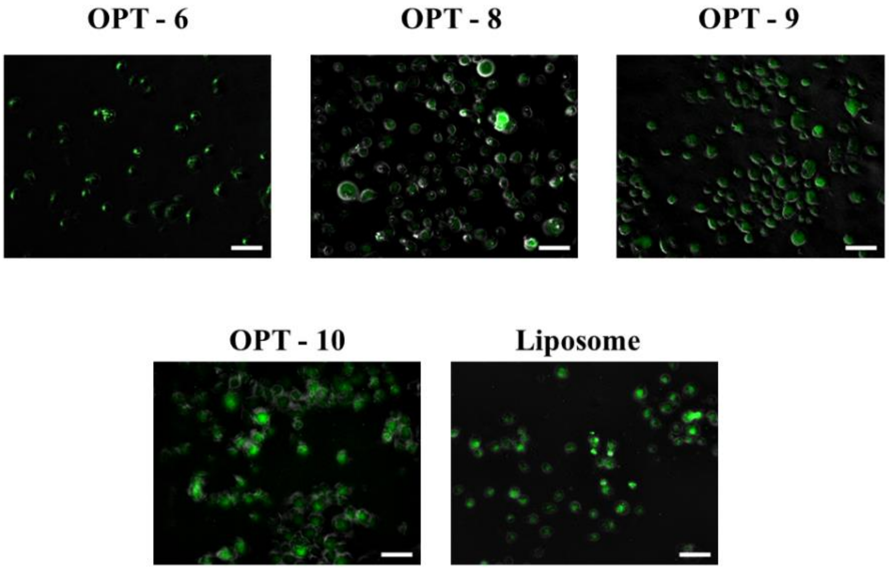


*Supplementary Figure 10: Confocal laser scanning microscopy analysis right after flow cytometry for the promising formulations. The same analysis should be repeated for the attached version of glioblastoma cells. White scaled label is presenting 1 µm.*

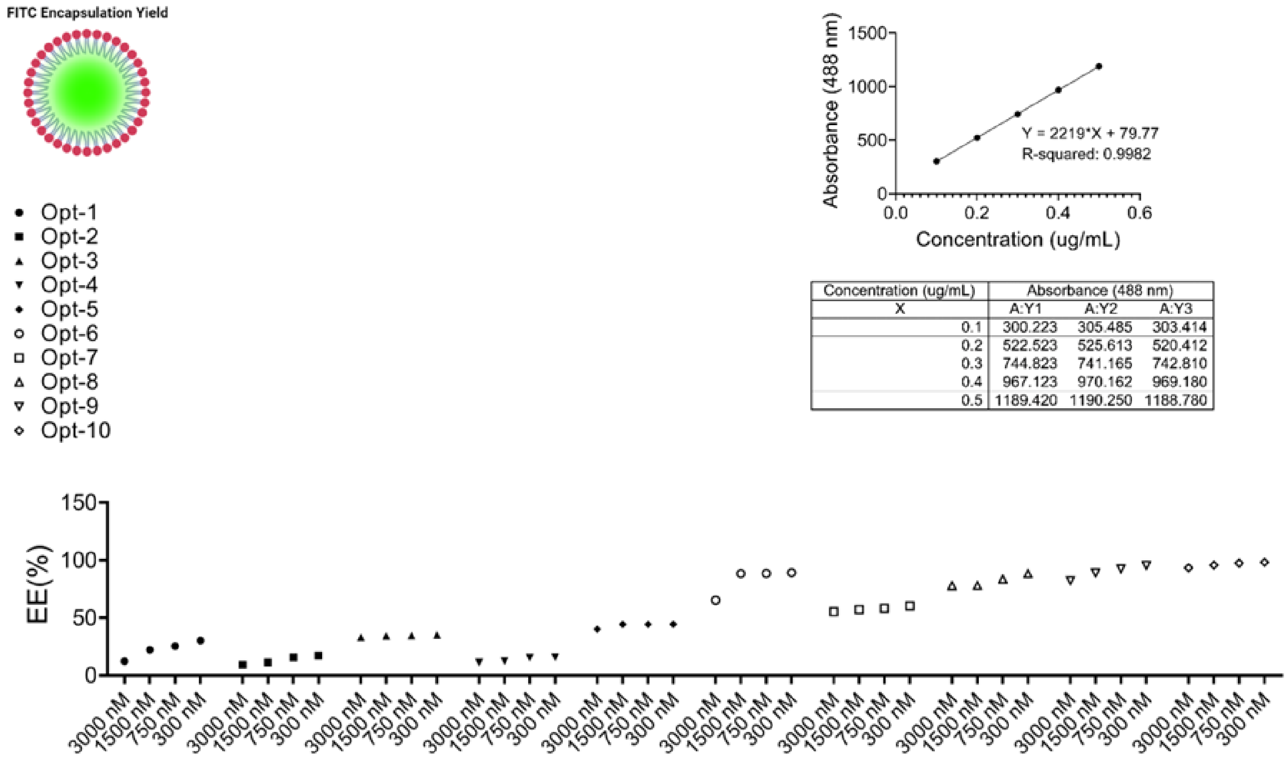


| Concentration (ug/mL) | Absorbance (488 nm) | | |
|---|---|---|---|
| X | A:Y1 | A:Y2 | A:Y3 |
| 0.1 | 300.223 | 305.485 | 303.414 |
| 0.2 | 522.523 | 525.613 | 520.412 |
| 0.3 | 744.823 | 741.165 | 742.810 |
| 0.4 | 967.123 | 970.162 | 969.180 |
| 0.5 | 1189.420 | 1190.250 | 1188.780 |

*Supplementary Figure 11. Encapsulation Efficiency (EE%) calculated in FITC-labeling. The entrapment is calculated by using FITC standard curve indicated at the right corner.*

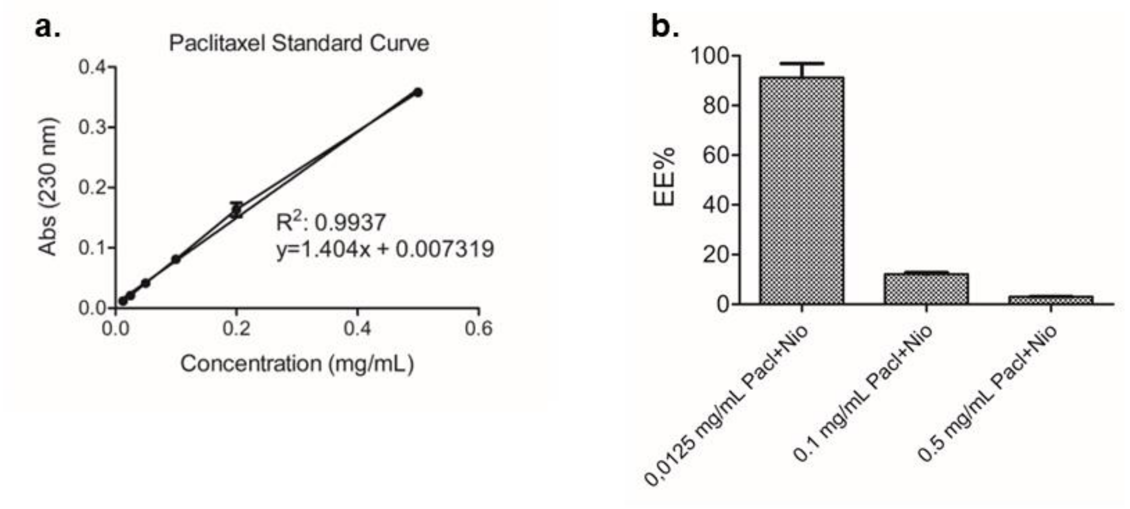


*Supplementary Figure 12. a. Standard curve for Paclitaxel obtained to calculate EE%. b. Encapsulation Efficiency (EE%) of three differently concentrated Paclitaxel-loaded niosomes.*

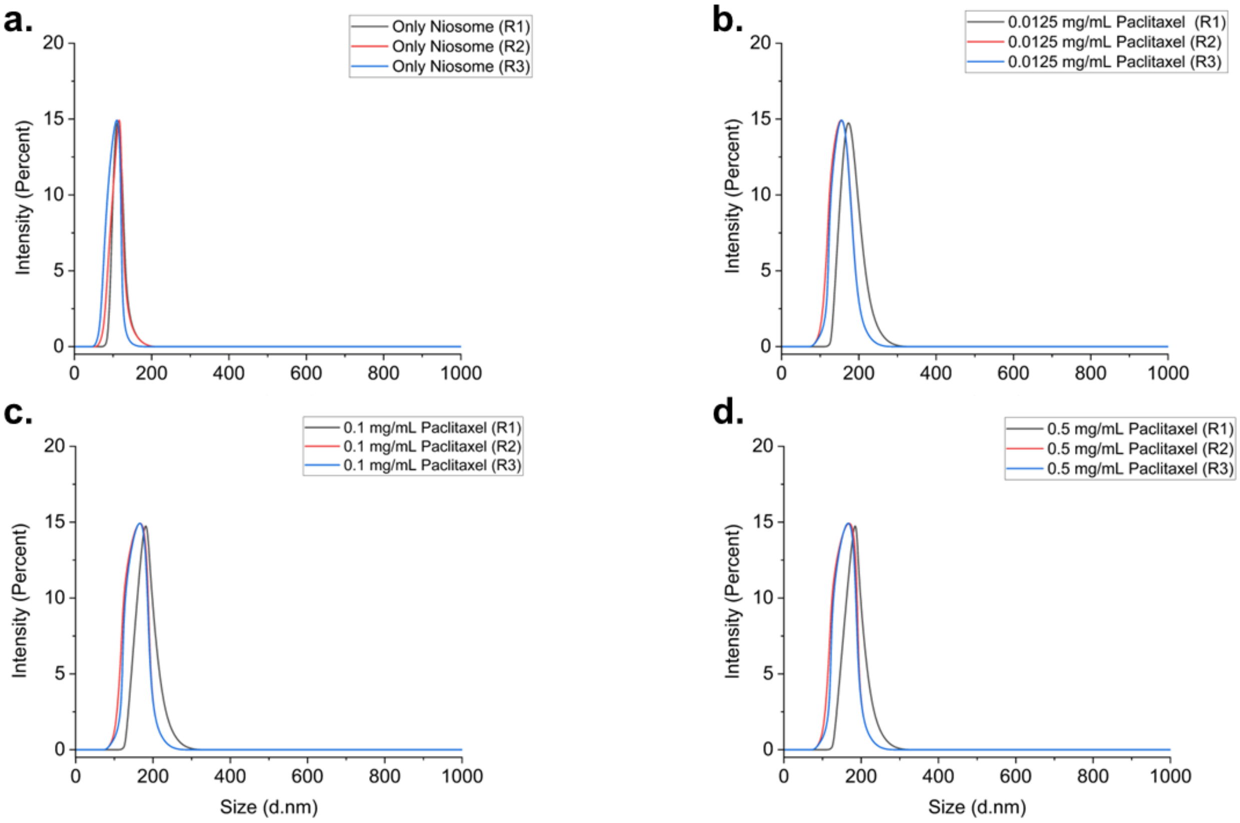


*Supplementary Figure 13. Dynamic Light Scattering (DLS) Gaussian distribution results for Paclitaxel-loaded niosomes. a. DLS results of only niosomes. b. 0.0125 mg/mL Paclitaxel-loaded niosomes. c. 0.1 mg/mL Paclitaxel-loaded niosomes. d. 0.5 mg/mL Paclitaxel-loaded niosomes.*

### Supplementary Tables

*Supplementary Table 1: Summary for effect analysis of sonication time by increased concentration for five different formulations – multiple non-paired t-test (non-parametric; non-paired)*

| Formulation A | Significance | P value | Mean of S.T*: 45 mins | Mean of S.T*: 90 mins | Difference | SE of difference | t ratio | df | q value |
|---|---|---|---|---|---|---|---|---|---|
| 0.1 mM | Yes | 0,000102 | 160,4 | 176,4 | -16 | 1,034 | 15,48 | 4 | 0,000103 |
| 0.2 mM | Yes | 0,000003 | 420,7 | 333,3 | 87,4 | 2,322 | 37,64 | 4 | 0,000004 |
| 0.3 mM | Yes | 0,000001 | 215,4 | 264,8 | -49,4 | 1,103 | 44,79 | 4 | 0,000003 |
| 0.42 mM | Yes | <0,000001 | 518,2 | 379,7 | 138,5 | 1,82 | 76,1 | 4 | <0,000001 |
| Formulation B | Significance | P value | Mean of S.T: 45 mins | Mean of S.T: 90 mins | Difference | SE of difference | t ratio | df | q value |
| 0.1 mM | Yes | 0,006845 | 107,3 | 114 | -6,7 | 1,306 | 5,129 | 4 | 0,007828 |
| 0.2 mM | Yes | 0,00008 | 126,7 | 120,5 | 6,2 | 0,3771 | 16,44 | 4 | 0,000243 |
| 0.3 mM | Yes | 0,007751 | 110,9 | 107,6 | 3,367 | 0,6799 | 4,952 | 4 | 0,007828 |
| 0.42 mM | No | 0,369881 | 103,7 | 106,2 | -2,567 | 2,543 | 1,009 | 4 | 0,280185 |
| Formulation C | Significance | P value | Mean of S.T: 45 mins | Mean of S.T: 90 mins | Difference | SE of difference | t ratio | df | q value |
| 0.1 mM | Yes | <0,000001 | 134,8 | 189,5 | -54,67 | 0,7149 | 76,47 | 4 | <0,000001 |
| 0.2 mM | Yes | 0,023326 | 150,1 | 152,2 | -2,1 | 0,5878 | 3,572 | 4 | 0,00589 |
| 0.3 mM | Yes | 0,000019 | 151,4 | 166,2 | -14,73 | 0,6236 | 23,63 | 4 | 0,000006 |
| 0.42 mM | Yes | <0,000001 | 172,1 | 194,4 | -22,3 | 0,2055 | 108,5 | 4 | <0,000001 |
| Formulation D | Significance | P value | Mean of S.T: 45 mins | Mean of S.T: 90 mins | Difference | SE of difference | t ratio | df | q value |
| 0.1 mM | Yes | 0,000003 | 129 | 103,3 | 25,7 | 0,6574 | 39,09 | 4 | 0,000003 |
| 0.2 mM | Yes | 0,000158 | 152,9 | 136,2 | 16,63 | 1,202 | 13,83 | 4 | 0,00008 |
| 0.3 mM | Yes | 0,002918 | 176,3 | 198,7 | -22,4 | 3,455 | 6,483 | 4 | 0,000983 |
| 0.42 mM | No | 0,992576 | 197,5 | 197,4 | 0,03333 | 3,367 | 0,009899 | 4 | 0,250625 |
| Formulation E | Significance | P value | Mean of S.T: 45 mins | Mean of S.T: 90 mins | Difference | SE of difference | t ratio | df | q value |
| 0.1 mM | No | 0,06341 | 193,5 | 206,1 | -12,63 | 4,957 | 2,548 | 4 | 0,032022 |
| 0.2 mM | Yes | 0,00003 | 222 | 271,2 | -49,2 | 2,334 | 21,08 | 4 | 0,00003 |
| 0.3 mM | Yes | 0,000012 | 282,6 | 190,2 | 92,37 | 3,491 | 26,46 | 4 | 0,000024 |
| 0.42 mM | No | 0,014917 | 211,4 | 198,4 | 13,03 | 3,183 | 4,095 | 4 | 0,010044 |

*Supplementary Table 2: Unpaired t-test statistical analysis results for concentration and sonication time parameters on pre-set criteria of size (100 to 200 nm); PDI (below 0.5), and zeta potential (-10 mV to 10 mV) through each preliminary niosome formulation. ns: p-value > 0.05; *: p-value ≤ 0.05; **: p-value ≤ 0.01; ***: p-value ≤ 0.001, ****: p-value ≤ 0.0001.*

Formulation Statistics

| FACTOR | Formulation ID | r | 95% Confidence Interval | R squared | P value (two-tailed) | Significant (alpha:0.05) | P value Summary |
|---|---|---|---|---|---|---|---|
| MOLARITY OF INGREDIENTS | Formulation A | 0,67 | 0,16 - 0,9 | 0,45 | 0,0157 | Yes | * |
| | Formulation B | -0,43 | -0,8 - 0,18 | 0,18 | 0,158 | No | NS |
| | Formulation C | 0,96 | 0,88 - 0,99 | 0,93 | < 0,0001 | Yes | **** |
| | Formulation D | 0,99 | 0,98 - 0,99 | 0,99 | < 0,0001 | Yes | **** |
| | Formulation E | 0,19 | -0,42 - 0,69 | 0,03 | 0,53 | No | NS |
| SONICATION TIME | Formulation A | -0,16 | 0,53 - 0,25 | 0,028 | 0,429 | No | NS |
| | Formulation B | -0,17 | 0,54 - 0,24 | 0,03 | 0,409 | No | NS |
| | Formulation C | 0,58 | 0,23 - 0,80 | 0,34 | 0,0026 | Yes | ** |
| | Formulation D | -0,02 | 0,42 - 0,37 | 0,0008 | 0,895 | No | NS |
| | Formulation E | -0,17 | -0,53 - 0,24 | 0,029 | 0,42 | No | NS |

*Supplementary Table 3: One-way ANOVA test for concentration parameter effect on PDI and zeta potential through each formulation for A. PDI, B. Zeta Potential. ns: p-value > 0.05; *: p-value ≤ 0.05; **: p-value ≤ 0.01; ***: p-value ≤ 0.001, ****: p-value ≤ 0.0001.*

| A | FACTOR | Two-way ANOVA multiple comparisons test | Mean Difference | 95,00% CI of Difference | Summary | Adjusted P Value | |
|---|---|---|---|---|---|---|---|
| | PDI | Formulation A vs. Formulation B | -0,09575 | -0,2644 to 0,07289 | ns | 0,4337 | A-B |
| | | Formulation A vs. Formulation C | 0,1235 | -0,04514 to 0,2921 | ns | 0,2109 | A-C |
| | | Formulation A vs. Formulation D | 0,01192 | -0,1567 to 0,1806 | ns | 0,9994 | A-D |
| | | Formulation A vs. Formulation E | 0,0545 | -0,1141 to 0,2231 | ns | 0,8522 | A-E |
| | | Formulation B vs. Formulation C | 0,2193 | 0,05061 to 0,3879 | ** | 0,0085 | B-C |
| | | Formulation B vs. Formulation D | 0,1077 | -0,06097 to 0,2763 | ns | 0,325 | B-D |
| | | Formulation B vs. Formulation E | 0,1503 | -0,01839 to 0,3189 | ns | 0,0924 | B-E |
| | | Formulation C vs. Formulation D | -0,1116 | -0,2802 to 0,05705 | ns | 0,2934 | C-D |
| | | Formulation C vs. Formulation E | -0,069 | -0,2376 to 0,09964 | ns | 0,7161 | C-E |
| | | Formulation D vs. Formulation E | 0,04258 | -0,1261 to 0,2112 | ns | 0,9326 | D-E |
| **B** | **FACTOR** | **Two-way ANOVA multiple comparisons test** | **Mean Difference** | **95,00% CI of difference** | **Summary** | **Adjusted P Value** | |
| | Zeta Potential (mV) | Formulation A vs. Formulation B | 0,1865 | -0,3072 to 0,6803 | ns | 0,7695 | A-B |
| | | Formulation A vs. Formulation C | -0,2286 | -0,7224 to 0,2651 | ns | 0,6193 | A-C |
| | | Formulation A vs. Formulation D | -0,004802 | -0,4986 to 0,4890 | ns | >0,9999 | A-D |
| | | Formulation A vs. Formulation E | 0,2515 | -0,2423 to 0,7453 | ns | 0,535 | A-E |
| | | Formulation B vs. Formulation C | -0,4152 | -0,9090 to 0,07860 | ns | 0,1212 | B-C |
| | | Formulation B vs. Formulation D | -0,1914 | -0,6851 to 0,3024 | ns | 0,7533 | B-D |
| | | Formulation B vs. Formulation E | 0,06498 | -0,4288 to 0,5588 | ns | 0,9936 | B-E |
| | | Formulation C vs. Formulation D | 0,2238 | -0,2700 to 0,7176 | ns | 0,637 | C-D |
| | | Formulation C vs. Formulation E | 0,4802 | -0,01362 to 0,9740 | ns | 0,0585 | C-E |
| | | Formulation D vs. Formulation E | 0,2563 | -0,2375 to 0,7501 | ns | 0,5176 | D-E |

*Supplementary Table 4A: Stability profile of optimized formulations in PDI for 21 days, 27 days, 35 days, and 92 days. Unpaired t-test is performed for each formulation. ns: p-value > 0.05; *: p-value ≤ 0.05; **: p-value ≤ 0.01; ***: p-value ≤ 0.001, ****: p-value ≤ 0.0001. Representative data is shown in Supplementary Fig 2.A.*

| FACTOR | Formulation ID | Actual mean | Number of values | t, df | P value (two tailed) | P value summary | Significant (alpha=0.05)? |
|---|---|---|---|---|---|---|---|
| PDI (1-21 days) | Opt-1 | 0,2263 | 2 | t=7,147, df=1 | 0,0885 | ns | No |
| | Opt-2 | 0,2438 | 2 | t=11,52, df=1 | 0,0551 | ns | No |
| | Opt-3 | 0,2982 | 2 | t=3,984, df=1 | 0,1565 | ns | No |
| | Opt-4 | 0,1678 | 2 | t=23,42, df=1 | 0,0272 | * | Yes |
| | Opt-5 | 0,2817 | 2 | t=281,7, df=1 | 0,0023 | ** | Yes |
| | Opt-6 | 0,1617 | 2 | t=3,212, df=1 | 0,1921 | ns | No |
| | Opt-7 | 0,02347 | 2 | t=1,903, df=1 | 0,3081 | ns | No |
| | Opt-8 | 0,04683 | 2 | t=1,222, df=1 | 0,4367 | ns | No |
| | Opt-9 | 0,1297 | 2 | t=7,939, df=1 | 0,0798 | ns | No |
| | Opt-10 | 0,1367 | 2 | t=12,81, df=1 | 0,0496 | * | Yes |
| PDI (1-27 Days) | Opt-1 | 0,2806 | 3 | t=4,903, df=2 | 0,0392 | * | Yes |
| | Opt-2 | 0,2559 | 3 | t=14,91, df=2 | 0,0045 | ** | Yes |
| | Opt-3 | 0,3626 | 3 | t=4,676, df=2 | 0,0428 | * | Yes |
| | Opt-4 | 0,149 | 3 | t=7,727, df=2 | 0,0163 | * | Yes |
| | Opt-5 | 0,275 | 3 | t=41,10, df=2 | 0,0006 | *** | Yes |
| | Opt-6 | 0,194 | 3 | t=4,463, df=2 | 0,0467 | * | Yes |
| | Opt-7 | 0,03464 | 3 | t=2,614, df=2 | 0,1205 | ns | No |
| | Opt-8 | 0,06011 | 3 | t=2,329, df=2 | 0,1452 | ns | No |
| | Opt-9 | 0,1164 | 3 | t=7,170, df=2 | 0,0189 | * | Yes |
| | Opt-10 | 0,1307 | 3 | t=15,20, df=2 | 0,0043 | ** | Yes |
| PDI (1-35 Days) | Opt-1 | 0,3009 | 4 | t=6,643, df=3 | 0,0069 | ** | Yes |
| | Opt-2 | 0,2838 | 4 | t=9,316, df=3 | 0,0026 | ** | Yes |
| | Opt-3 | 0,4375 | 4 | t=4,711, df=3 | 0,0181 | * | Yes |
| | Opt-4 | 0,166 | 4 | t=7,617, df=3 | 0,0047 | ** | Yes |
| | Opt-5 | 0,2703 | 4 | t=40,68, df=3 | <0,0001 | **** | Yes |
| | Opt-6 | 0,2319 | 4 | t=4,751, df=3 | 0,0177 | * | Yes |
| | Opt-7 | 0,0679 | 4 | t=1,965, df=3 | 0,1441 | ns | No |
| | Opt-8 | 0,1007 | 4 | t=2,264, df=3 | 0,1086 | ns | No |
| | Opt-9 | 0,1123 | 4 | t=9,181, df=3 | 0,0027 | ** | Yes |
| | Opt-10 | 0,1157 | 4 | t=7,146, df=3 | 0,0056 | ** | Yes |
| PDI (1-92 Days) | Opt-1 | 0,3498 | 5 | t=5,813, df=4 | 0,0044 | ** | Yes |
| | Opt-2 | 0,2403 | 5 | t=4,849, df=4 | 0,0083 | ** | Yes |
| | Opt-3 | 0,4005 | 5 | t=4,950, df=4 | 0,0078 | ** | Yes |
| | Opt-4 | 0,161 | 5 | t=9,145, df=4 | 0,0008 | *** | Yes |
| | Opt-5 | 0,2665 | 5 | t=41,66, df=4 | <0,0001 | **** | Yes |
| | Opt-6 | 0,2773 | 5 | t=4,696, df=4 | 0,0093 | ** | Yes |
| | Opt-7 | 0,1071 | 5 | t=2,256, df=4 | 0,0871 | ns | No |
| | Opt-8 | 0,1101 | 5 | t=3,083, df=4 | 0,0368 | * | Yes |
| | Opt-9 | 0,1264 | 5 | t=7,424, df=4 | 0,0018 | ** | Yes |
| | Opt-10 | 0,1271 | 5 | t=7,483, df=4 | 0,0017 | ** | Yes |

*Supplementary Table 4B: Stability profile of optimized formulations in Zeta Potential for 21 days, 27 days, 35 days, and 92 days. Unpaired t-test is performed for each formulation. ns: p-value > 0.05; *: p-value ≤ 0.05; **: p-value ≤ 0.01; ***: p-value ≤ 0.001, ****: p-value ≤ 0.0001. Representative data is shown in Supplementary Fig 2.B.*

| FACTOR | Formulation ID | Actual mean | Number of values | t, df | P value (two tailed) | P value summary | Significant (alpha=0.05)? |
|---|---|---|---|---|---|---|---|
| Zeta Potential (1-21 days) | Opt-1 | -0,00541 | 2 | t=1,308, df=1 | 0,4156 | ns | No |
| | Opt-2 | -0,5948 | 2 | t=12,52, df=1 | 0,0507 | ns | No |
| | Opt-3 | 0,2827 | 2 | t=5,616, df=1 | 0,1122 | ns | No |
| | Opt-4 | -0,1742 | 2 | t=26,79, df=1 | 0,0237 | * | Yes |
| | Opt-5 | -0,4862 | 2 | t=5,535, df=1 | 0,1138 | ns | No |
| | Opt-6 | -0,4765 | 2 | t=8,287, df=1 | 0,0765 | ns | No |
| | Opt-7 | -0,6827 | 2 | t=17,07, df=1 | 0,0373 | * | Yes |
| | Opt-8 | -0,3832 | 2 | t=7,074, df=1 | 0,0894 | ns | No |
| | Opt-9 | -0,0476 | 2 | t=1,372, df=1 | 0,401 | ns | No |
| | Opt-10 | -0,253 | 2 | t=126,5, df=1 | 0,005 | ** | Yes |
| Zeta Potential (1-27 days) | Opt-1 | -0,0605 | 3 | t=1,097, df=2 | 0,387 | ns | No |
| | Opt-2 | -0,4115 | 3 | t=2,219, df=2 | 0,1567 | ns | No |
| | Opt-3 | 0,1897 | 3 | t=1,949, df=2 | 0,1906 | ns | No |
| | Opt-4 | -0,1663 | 3 | t=19,15, df=2 | 0,0027 | ** | Yes |
| | Opt-5 | -0,4514 | 3 | t=7,345, df=2 | 0,018 | * | Yes |
| | Opt-6 | -0,4423 | 3 | t=9,285, df=2 | 0,0114 | * | Yes |
| | Opt-7 | -0,5694 | 3 | t=4,928, df=2 | 0,0388 | * | Yes |
| | Opt-8 | -0,4672 | 3 | t=5,210, df=2 | 0,0349 | * | Yes |
| | Opt-9 | -0,08248 | 3 | t=2,051, df=2 | 0,1768 | ns | No |
| | Opt-10 | -0,2438 | 3 | t=26,23, df=2 | 0,0015 | ** | Yes |
| Zeta Potential (1-35 days) | Opt-1 | -0,1752 | 4 | t=1,446, df=3 | 0,2439 | ns | No |
| | Opt-2 | -0,3523 | 4 | t=2,449, df=3 | 0,0918 | ns | No |
| | Opt-3 | 0,04806 | 4 | t=0,3051, df=3 | 0,7802 | ns | No |
| | Opt-4 | -0,3215 | 4 | t=2,070, df=3 | 0,1302 | ns | No |
| | Opt-5 | -0,4343 | 4 | t=9,292, df=3 | 0,0026 | ** | Yes |
| | Opt-6 | -0,426 | 4 | t=11,38, df=3 | 0,0015 | ** | Yes |
| | Opt-7 | -0,6333 | 4 | t=6,106, df=3 | 0,0088 | ** | Yes |
| | Opt-8 | -0,4413 | 4 | t=6,439, df=3 | 0,0076 | ** | Yes |
| | Opt-9 | -0,1438 | 4 | t=2,128, df=3 | 0,1233 | ns | No |
| | Opt-10 | -0,2458 | 4 | t=35,82, df=3 | <0,0001 | **** | Yes |
| Zeta Potential (1-92 days) | Opt-1 | -0,144 | 5 | t=1,456, df=4 | 0,219 | ns | No |
| | Opt-2 | -0,2699 | 5 | t=1,948, df=4 | 0,1232 | ns | No |
| | Opt-3 | 0,08311 | 5 | t=0,6547, df=4 | 0,5484 | ns | No |
| | Opt-4 | -0,3137 | 5 | t=2,603, df=4 | 0,0599 | ns | No |
| | Opt-5 | -0,3671 | 5 | t=4,814, df=4 | 0,0086 | ** | Yes |
| | Opt-6 | -0,3804 | 5 | t=7,039, df=4 | 0,0021 | ** | Yes |
| | Opt-7 | -0,6163 | 5 | t=7,505, df=4 | 0,0017 | ** | Yes |
| | Opt-8 | -0,4674 | 5 | t=7,899, df=4 | 0,0014 | ** | Yes |
| | Opt-9 | -0,2713 | 5 | t=1,968, df=4 | 0,1204 | ns | No |
| | Opt-10 | -0,3939 | 5 | t=2,657, df=4 | 0,0566 | ns | No |

*Supplementary Table 5: Reproducibility profile of optimized formulations in size (nm), PDI, and zeta potential (mV). Unpaired t-test is performed for each formulation. ns: p-value > 0.05; *: p-value ≤ 0.05; **: p-value ≤ 0.01; ***: p-value ≤ 0.001, ****: p-value ≤ 0.0001.*

| FACTOR | Formulation ID | Actual mean | Number of values | t, df | P value (two tailed) | P value summary | Significant (alpha=0.05)? |
|---|---|---|---|---|---|---|---|
| Size (nm) | Opt-1 | 112.4 | 3 | t=7,389, df=2 | 0,0178 | * | Yes |
| | Opt-2 | 122.3 | 3 | t=35,80, df=2 | 0,0008 | ns | No |
| | Opt-3 | 120 | 3 | t=18,77, df=2 | 0,0028 | ** | Yes |
| | Opt-4 | 160.3 | 3 | t=19,15, df=2 | 0,0027 | ** | Yes |
| | Opt-5 | 219.6 | 3 | t=6,154, df=2 | 0,0254 | * | Yes |
| | Opt-6 | 109.6 | 3 | t=8,108, df=2 | 0,0149 | * | Yes |
| | Opt-7 | 242.9 | 3 | t=7,827, df=2 | 0,0159 | * | Yes |
| | Opt-8 | 169.5 | 3 | t=10,40, df=2 | 0,0091 | ** | Yes |
| | Opt-9 | 170.2 | 3 | t=9,159, df=2 | 0,0117 | ns | No |
| | Opt-10 | 154.5 | 3 | t=74,75, df=2 | 0,0002 | ns | No |
| PDI | Opt-1 | 0,317 | 3 | t=8,115, df=2 | 0,0148 | * | Yes |
| | Opt-2 | 0,2684 | 3 | t=10,88, df=2 | 0,0083 | ** | Yes |
| | Opt-3 | 0,3727 | 3 | t=4,633, df=2 | 0,0436 | * | Yes |
| | Opt-4 | 0,1504 | 3 | t=6,325, df=2 | 0,0241 | * | Yes |
| | Opt-5 | 0,2748 | 3 | t=26,31, df=2 | 0,0014 | ** | Yes |
| | Opt-6 | 0,1932 | 3 | t=74,18, df=2 | 0,0002 | *** | Yes |
| | Opt-7 | 0,04592 | 3 | t=2,708, df=2 | 0,1136 | ns | No |
| | Opt-8 | 0,07678 | 3 | t=17,98, df=2 | 0,0031 | ** | Yes |
| | Opt-9 | 0,08767 | 3 | t=11,81, df=2 | 0,0071 | ns | No |
| | Opt-10 | 0,1288 | 3 | t=18,47, df=2 | 0,0029 | ns | No |
| Zeta Potential (mV) | Opt-1 | -0,06303 | 3 | t=1,154, df=2 | 0,3679 | ns | No |
| | Opt-2 | -0,05773 | 3 | t=0,8288,df=2 | 0,4944 | ns | No |
| | Opt-3 | 0,1985 | 3 | t=1,973, df=2 | 0,1872 | ns | No |
| | Opt-4 | -0,1699 | 3 | t=20,89, df=2 | 0,0023 | ** | Yes |
| | Opt-5 | -0,4786 | 3 | t=8,751, df=2 | 0,0128 | * | Yes |
| | Opt-6 | -0,4493 | 3 | t=6,825, df=2 | 0,0208 | * | * |
| | Opt-7 | -0,4636 | 3 | t=9,101, df=2 | 0,0119 | * | Yes |
| | Opt-8 | -0,5732 | 3 | t=5,254, df=2 | 0,0344 | * | Yes |
| | Opt-9 | -0,08619 | 3 | t=2,030, df=2 | 0,1795 | ns | No |
| | Opt-10 | -0,2558 | 3 | t=31,57, df=2 | 0,001 | ns | No |

*Supplementary Table 6: Dynamic Light Scattering data for Liposome*

| **Liposome (Dye (-))** | **Measurements** | | | **Mean** | **SD** |
|---|---|---|---|---|---|
| | 1 | 2 | 3 | | |
| Size (nm) | 179,7 | 185,5 | 183,3 | 182,83 | 2,93 |
| PDI | 0,144 | 0,102 | 0,138 | 0,13 | 0,02 |
| Zeta Potential (mV) | -6,51 | -6,35 | -6,48 | -6,45 | 0,09 |
| **Liposome (Dye (+))** | **Measurements** | | | **Mean** | **SD** |
| | 1 | 2 | 3 | | |
| Size (nm) | 192,11 | 189,13 | 201,33 | 194,190 | 6,360 |
| PDI | 0,266 | 0,307 | 0,283 | 0,285 | 0,021 |
| Zeta Potential (mV) | -0,122 | -0,352 | 0,144 | -0,110 | 0,248 |

*Supplementary Table 7. Previously loaded FITC concentration versus obtained absorbance, released FITC concentration based on received FITC absorbance, calculated and EE% for each niosome formulation.*

| | | | Measured Absorbance (a.u.) | | | |
|---|---|---|---|---|---|---|
| **Percentage (v/v%)** | Sample ID | Loaded Concentration (nM) | R1 | R2 | R3 | Calculated EE% |
| **0,1** | Opt-1 | 3000 | 819,78 | 821,1 | 818,22 | 12,3 |
| **0,05** | | 1500 | 740,69 | 751,5 | 746,77 | 22,2 |
| **0,025** | | 750 | 442,98 | 455,1 | 444,21 | 25,4 |
| **0,01** | | 300 | 225,13 | 235,4 | 244,36 | 30,2 |
| **0,1** | Opt-2 | 3000 | 620,45 | 623,4 | 633,43 | 9,21 |
| **0,05** | | 1500 | 382,74 | 388,8 | 377,89 | 11,23 |
| **0,025** | | 750 | 273,11 | 287,1 | 281,22 | 15,66 |
| **0,01** | | 300 | 127,54 | 133,3 | 128,73 | 17,11 |
| **0,1** | Opt-3 | 3000 | 2236,6 | 2310 | 2214,56 | 33,2 |
| **0,05** | | 1500 | 2313,4 | 2345 | 2244,2 | 34,34 |
| **0,025** | | 750 | 2378,22 | 2406 | 2403,66 | 34,56 |
| **0,01** | | 300 | 2368,33 | 2401 | 2389,77 | 35,16 |
| **0,1** | Opt-4 | 3000 | 757,88 | 784,3 | 765,46 | 11,25 |
| **0,05** | | 1500 | 420,23 | 422,3 | 425,66 | 12,33 |
| **0,025** | | 750 | 269,27 | 278 | 288,12 | 15,44 |
| **0,01** | | 300 | 117,56 | 119,2 | 118,23 | 15,77 |
| **0,1** | Opt-5 | 3000 | 2701,44 | 2677 | 2544,33 | 40,1 |
| **0,05** | | 1500 | 1506,8 | 1510 | 1522,12 | 44,21 |
| **0,025** | | 750 | 771,03 | 782,3 | 777,34 | 44,23 |
| **0,01** | | 300 | 329,77 | 333,5 | 341,45 | 44,35 |
| **0,1** | Opt-6 | 3000 | 4399,11 | 4223 | 4456,23 | 65,3 |
| **0,05** | | 1500 | 3010,86 | 3112 | 3078,22 | 88,34 |
| **0,025** | | 750 | 1472,03 | 1488 | 1477,12 | 88,45 |
| **0,01** | | 300 | 593,27 | 599,2 | 598,22 | 89,12 |
| **0,1** | Opt-7 | 3000 | 3674,66 | 3702 | 3508,22 | 55,2 |
| **0,05** | | 1500 | 1904,901 | 1920 | 1911,2 | 57,23 |
| **0,025** | | 750 | 970,75 | 966,3 | 981,23 | 58,33 |
| **0,01** | | 300 | 401,61 | 410,2 | 405,67 | 60,33 |
| **0,1** | Opt-8 | 3000 | 5184,47 | 5089 | 5088,12 | 77,88 |
| **0,05** | | 1500 | 2600,22 | 2712 | 2683,22 | 78,12 |
| **0,025** | | 750 | 1300,11 | 1322 | 1311,44 | 83,45 |
| **0,01** | | 300 | 520,44 | 522,3 | 530,24 | 88,48 |
| **0,1** | Opt-9 | 3000 | 5566,06 | 5603 | 5449,04 | 82,11 |
| **0,05** | | 1500 | 2960,03 | 2888 | 2902,33 | 88,93 |
| **0,025** | | 750 | 1538,6 | 1522 | 1507,45 | 92,45 |
| **0,01** | | 300 | 635,47 | 644,3 | 645,11 | 95,46 |
| **0,1** | Opt-10 | 3000 | 6220,31 | 6122 | 6114,66 | 93,44 |
| **0,05** | | 1500 | 3167,23 | 3056 | 3077,88 | 95,67 |
| **0,025** | | 750 | 1619,81 | 1607 | 1623,22 | 97,33 |
| **0,01** | | 300 | 653,91 | 666,2 | 672,34 | 98,23 |

*Supplementary Table 8. Encapsulation Efficiency (EE%), Drug Loading Capacity (DLC%), and Drug Loading Efficiency (DLE%) of FITC-loaded niosomes determined for 300 nM FITC with 0.5% of total volumetric ratio.*

| **Niosomes** | **EE%** | **DLC%** | **DLE%** |
|---|---|---|---|
| Opt-1 | 42.1 ± 3.2 | 4.7 ± 2.5 | 34.4 ± 0.7 |
| Opt-2 | 23.5 ± 1.7 | 2.1 ± 0.2 | 14.2 ± 1.2 |
| Opt-3 | 35.2 ± 2.6 | 2.5 ± 1.1 | 21.2 ± 1.4 |
| Opt-4 | 12.4 ± 0.5 | 0.93 ± 0.5 | 9.1 ± 0.3 |
| Opt-5 | 52.3 ± 0.3 | 4.8 ± 0.8 | 44.2 ± 2.2 |
| Opt-6 | 92.1 ± 2.2 | 8.9 ± 3.1 | 87.3 ± 3.5 |
| Opt-7 | 57.6 ± 4.1 | 5.4 ± 2.2 | 48.3 ± 2.1 |
| Opt-8 | 85.4 ± 1.8 | 7.7 ± 0.5 | 77.2 ± 3.1 |
| Opt-9 | 95.1 ± 0.2 | 9.01 ± 0.7 | 88.4 ± 0.3 |
| Opt-10 | 96.2 ± 1.6 | 9.4 ± 1.4 | 90.9 ± 0.9 |
